# Niche-specific profiling reveals transcriptional adaptations required for the cytosolic lifestyle of *Salmonella enterica*

**DOI:** 10.1101/2021.01.11.426201

**Authors:** TuShun R. Powers, Amanda L. Haeberle, Disa L. Hammarlöf, Jennifer A. Cundiff, Alexander V. Predeus, Karsten Hokamp, Jay C.D. Hinton, Leigh A. Knodler

## Abstract

*Salmonella enterica* serovar Typhimurium (*S*. Typhimurium) is a zoonotic pathogen that causes diarrheal disease in humans and animals. During salmonellosis, *S*. Typhimurium colonizes epithelial cells lining the gastrointestinal tract. *S*. Typhimurium has an unusual lifestyle in epithelial cells that begins within an endocytic-derived *Salmonella*-containing vacuole (SCV), followed by escape into the cytosol, epithelial cell lysis and bacterial release. The cytosol is a more permissive environment than the SCV and supports rapid bacterial growth. The physicochemical conditions encountered by *S.* Typhimurium within the cytosol, and the bacterial genes required for cytosolic colonization, remain unknown. Here we have exploited the parallel colonization strategies of *S*. Typhimurium in epithelial cells to decipher the two niche-specific bacterial virulence programs. By combining a population-based RNA-seq approach with single-cell microscopic analysis, we identified bacterial genes/sRNAs with cytosol-specific or vacuole-specific expression signatures. Using these genes/sRNAs as environmental biosensors, we defined that *Salmonella* is exposed to iron and manganese deprivation and oxidative stress in the cytosol and zinc and magnesium deprivation in the SCV. Furthermore, iron availability was critical for optimal *S*. Typhimurium replication in the cytosol, as well as *entC*, *fepB*, *soxS* and *sitA-mntH*. Virulence genes that are typically associated with extracellular bacteria, namely *Salmonella* pathogenicity island 1 (SPI1) and SPI4, had a cytosolic-specific expression profile. Our study reveals that the cytosolic and vacuolar *S*. Typhimurium virulence gene programs are unique to, and tailored for, residence within distinct intracellular compartments. Therefore, this archetypical vacuole-adapted pathogen requires extensive transcriptional reprogramming to successfully colonize the mammalian cytosol.

**Author Summary:** Intracellular pathogens reside either within a membrane-bound vacuole or are free-living in the cytosol and their virulence programs are tailored towards survival within a particular intracellular compartment. Some bacterial pathogens (such as *Salmonella enterica*) can successfully colonize both intracellular niches, but how they do so is unclear. Here we have exploited the parallel intracellular lifestyles of *S. enterica* in epithelial cells to identify the niche-specific bacterial expression profiles and environmental cues encountered by *S. enterica*. We have also discovered bacterial genes that are required for colonization of the cytosol, but not the vacuole. Our results advance our understanding of pathogen-adaptation to alternative replication niches and highlight an emerging concept in the field of bacteria-host cell interactions.

## Introduction

There are two major niches in which intracellular bacteria survive and proliferate after internalization into host cells, confined within a membrane-bound vacuole or free-living within the cytosol. Different pathogenic mechanisms are required to occupy these diverse environments. Historically, bacterial pathogens have been categorized as being either vacuolar or cytosolic, but it has recently been realized that some bacteria can occupy both niches, often in a cell-type specific manner. Examples are *Salmonella enterica*, *Mycobacterium tuberculosis* and *Listeria monocytogenes* [1–3]. What is unclear is how bacteria that are adapted to survive within one intracellular niche can successfully colonize a distinct cellular compartment.

Of the foodborne bacterial, protozoal and viral diseases, non-typhoidal *Salmonella enterica* (NTS) cause the largest burden of illness and death worldwide [4]. Infection can cause either a self-limiting gastroenteritis or a life-threatening, invasive disease (invasive non-typhoidal salmonellosis) in immunocompromised individuals, which is particularly a public health problem in sub-Saharan Africa and south-east Asia [5]. Upon ingestion of contaminated food or water, *S*. *enterica* enters epithelial cells lining the gut and resides within a membrane-bound compartment derived from the endocytic pathway [6, 7], the *Salmonella*-containing vacuole (SCV). Entry into non-phagocytic cells is largely governed by a type III secretion system, T3SS1, that is encoded by *Salmonella* pathogenicity island (SPI) 1. The T3SS acts as a molecular syringe to inject bacterial effector proteins across the eukaryotic cell plasma membrane to modulate host signaling networks that induce rearrangement of the actin cytoskeleton, leading to the formation of plasma membrane “ruffles” and engulfment of bacteria (reviewed in [8]). Establishment and maintenance of the SCV is dependent upon effectors being translocated across the vacuolar membrane by T3SS1 and another T3SS, T3SS2, which is specifically induced by environmental cues sensed by bacteria within the SCV lumen [9, 10]. T3SS2 is also important for survival within phagocytic cells, which *S*. *enterica* encounters in the lamina propria during an enteric infection or in the mesenteric lymph nodes, reticuloendothelial tissues (liver and spleen) and circulating blood during invasive disease.

*S*. *enterica* resides within a membrane-bound vacuole within epithelial cells, fibroblasts, macrophages and dendritic cells (reviewed in [11]). However, the intracellular lifestyle of *S. enterica* differs between cell types, with a proportion of the total bacterial population living freely in the cytosol of epithelial cells, a phenomenon described for *Salmonella enterica* serovar Typhimurium (*S*. Typhimurium) and *S*. Typhi infections *in vitro* and/or *in vivo* [12–16]. The eventual outcome of epithelial cells harboring cytosolic bacteria is their expulsion from the monolayer into the lumen of the gut and gall bladder [12, 17]; we have proposed that this is a mechanism utilized by *Salmonella* for spreading within and between hosts. Notably, the cytosol of macrophages is not permissive for *S.* Typhimurium replication [18, 19], possibly due to nutrient limitation or enhanced host cell innate immune defenses such as autophagy and/or inflammasomes [19]. Considering that the site of intracellular replication is cell-type restricted, we propose that *S. enterica* is an “opportunistic” cytosolic pathogen.

A specialized form of autophagy, called xenophagy, protects the mammalian cytosol by targeting intracellular pathogens to autophagosomes for their eventual degradation in lysosomes (reviewed in [20]). In fact, many of the components of the autophagic machinery have been identified using *S*. Typhimurium as a model pathogen. In the first report of autophagic recognition, Brumell and colleagues showed that a sub-population of internalized *S*. Typhimurium damage their nascent vacuole in epithelial cells in a T3SS1-dependent manner and these bacteria are decorated with the autophagy marker, microtubule-associated light chain-3 (LC3) [21]. A recently identified type III effector, SopF, limits the decoration of SCVs with LC3 [22, 23]. Damage exposes host glycans restricted to the vacuole lumen to the cytosol, which are then recognized by β-galactoside-binding lectins, specifically galectin-3 (GAL3), GAL8 and GAL9 [24]. Galectin binding acts as an “eat me” signal that initiates a cascade of receptor binding, phagophore formation and tethering to the bacterium, culminating in autophagosome formation [24, 25]. However, autophagic control of *Salmonella* is temporal and incomplete, at least in epithelial cells [12, 21]. Furthermore, *S*. Typhimurium can also use autophagy to repair damaged SCVs in mouse embryonic fibroblasts [26] and promote replication in the cytosol of epithelial cells [27]. Autophagy therefore serves both a pro- and anti-bacterial role in *S. enterica* infections.

Previous studies have largely investigated the infectious cycle of *S*. Typhimurium in epithelial cells by population-based analyses, which do not account for the heterogeneous population of intracellular bacteria. Only by determining the distinct responses of intracellular *Salmonella* to the cytosolic and the vacuolar niche can the infection biology of this important pathogen be accurately defined. Whilst the distinct milieus encountered within a vacuole versus the cytosol provide site-specific cues for *Salmonella* gene induction, little is known about what these cues might be, or which bacterial genes drive replication/survival in the cytosol. Using a combination of RNA-seq-based transcriptomics and single-cell microscopic analysis, here we define the niche-specific environments encountered by *S*. Typhimurium in epithelial cells and identify *S*. Typhimurium genes that are required for bacterial proliferation in the cytosol.

## Results

### Modulation of autophagy affects bacterial proliferation in the epithelial cell cytosol

We hypothesized that the modulation of autophagy in epithelial cells would selectively perturb the cytosolic, and not vacuolar, proliferation of *S*. Typhimurium. Autophagy levels can be manipulated by pharmacological or genetic means (reviewed in [28]). To test our hypothesis, we used nutrient starvation to upregulate autophagy and the class III phosphoinositide 3-kinase (PI3K) inhibitor, wortmannin (WTM), to inhibit autophagy (Fig 1A). To enumerate cytosolic bacteria after autophagy activation/inhibition, we used the digitonin permeabilization assay [12, 13] to label the bacteria accessible to cytosol-delivered anti-*S*. Typhimurium lipopolysaccharide (LPS) antibodies at the early stages of infection (15 min – 3 h p.i., Fig 1B). Wild-type bacteria were constitutively expressing *mCherry* from a plasmid, pFPV-mCherry. In untreated cells (basal levels of autophagy), 6.8% of bacteria were accessible to the cytosol as early as 15 min p.i. This proportion increased to ∼20% by 45 min p.i. and remained at a steady-state thereafter (Fig 1B). Upon activation of autophagy (EBSS), the fraction of bacteria that were accessible to the cytosol was reduced at 45 min, 90 min and 180 min p.i. (Fig 1B), consistent with enhanced autophagic capture of bacteria in, or repair of, damaged vacuoles. In contrast, inhibition of autophagy (WTM) increased the proportion of cytosolic bacteria at all timepoints examined (Fig 1B), in agreement with previous findings [14, 29].

**Fig 1.**
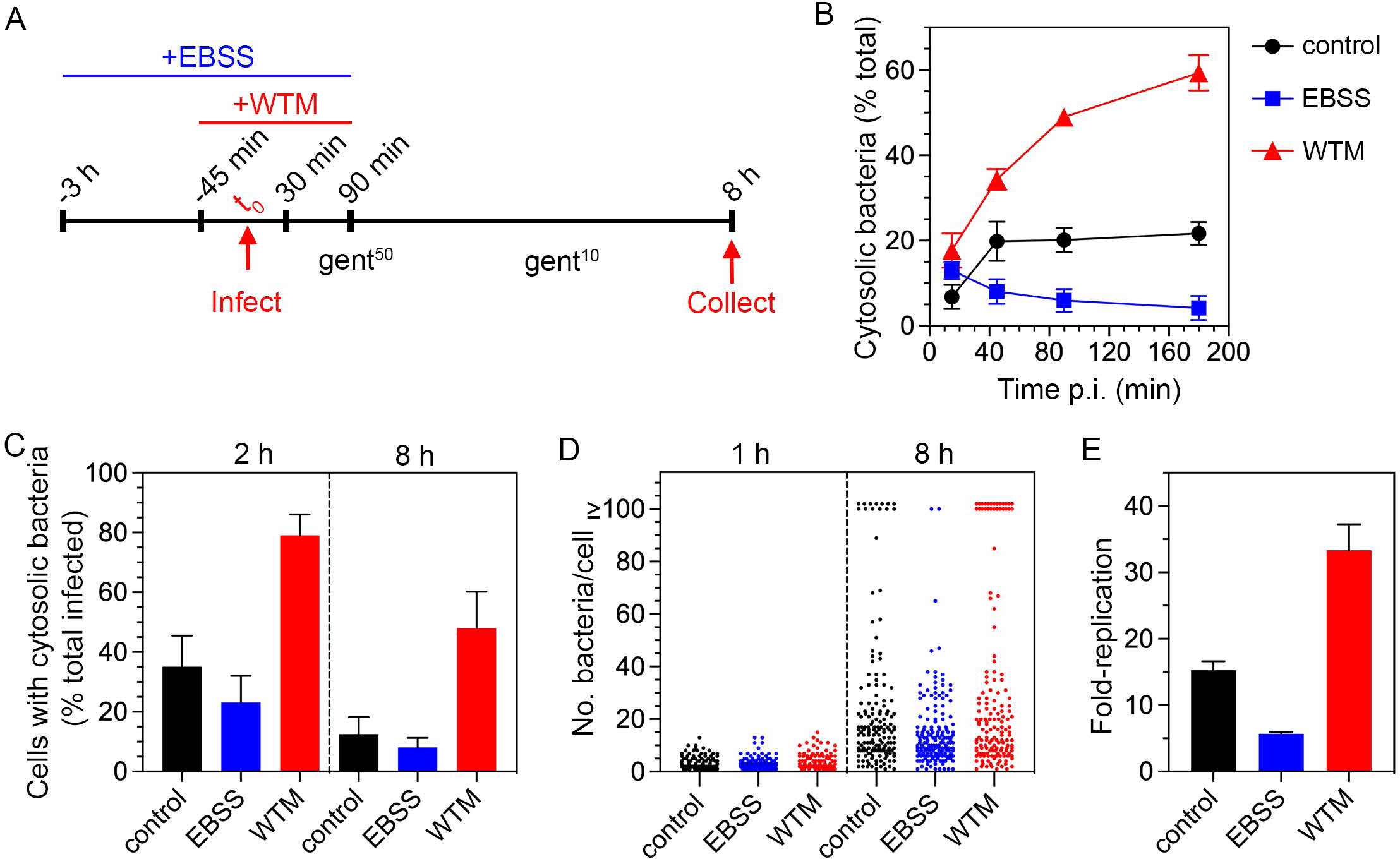
EBSS and wortmannin inversely affect the cytosolic population. (A) Schematic of the experimental design employed to differentially modulate host cell autophagy. (B) Epithelial cells seeded on glass coverslips were pretreated with EBSS or 100 nM WTM as depicted in (A) and infected with *S*. Typhimurium wild-type bacteria harboring pFPV-mCherry for plasmid-borne, constitutive expression of *mCherry*. The proportion of bacteria accessible to anti-*Salmonella* LPS antibodies delivered to the mammalian cell cytosol was determined by digitonin permeabilization assay and fluorescence microscopy. n≥3 independent experiments (C) Epithelial cells seeded on coverslips were pretreated as in (A) and infected with mCherry-S. Typhimurium (wild-type *glmS*::P*trc*-*mCherryST*::FRT bacteria constitutively expressing a chromosomal copy of *mCherry*) harboring the fluorescent reporter plasmid, P*uhpT*-*gfpova* (pNF101). At the indicated times, cells were fixed and the proportion of infected cells containing cytosolic bacteria (GFP-positive) was scored by fluorescence microscopy. n=3 independent experiments. (D) Cells seeded on coverslips were pretreated as in (A) and infected with wild-type bacteria harboring pFPV-mCherry. The number of bacteria in each infected cell was scored by fluorescence microscopy. Cells with ≥100 bacteria contain cytosolic *S*. Typhimurium. Each dot represents one infected cell. n=2 (1 h) or 3 (8 h) independent experiments. (E) Epithelial cells were infected with wild-type bacteria and the number of intracellular bacteria at 1 h and 8 h p.i. was determined by gentamicin protection assay. Fold-replication is CFUs at 8 h/1 h. n=3 independent experiments. For all panels, control = untreated cells; EBSS = Earle’s balanced salt solution treatment; WTM = wortmannin treatment.

To verify cytosolic localization, we used a transcriptional reporter plasmid, pNF101, that only expresses *gfp-ova* in the sub-population of *S*. Typhimurium that are in damaged vacuoles and/or free in the cytosol. Epithelial cells were infected with mCherry-*S*. Typhimurium (*S*. Typhimurium wild-type bacteria constitutively expressing *mCherry* on the chromosome) harboring pNF101. In untreated cells, 35% and 13% of infected cells harbored GFP-positive (cytosolic) bacteria at 2 h and 8 h p.i., respectively (Fig 1C). EBSS treatment lowered this proportion at both timepoints to 23% and 8%, respectively. Conversely, WTM treatment increased the level of infected cells with cytosolic bacteria at both 2 h and 8 h p.i. (79% and 48%).

To quantify the niche-specific effects of EBSS and WTM treatments on bacterial replication, epithelial cells were infected with *S*. Typhimurium harboring pFPV-mCherry and the number of bacteria per cell was scored. Cytosolic *S*. Typhimurium that evade autophagic clearance replicate at a much faster rate than vacuolar bacteria in epithelial cells [12,13,30], eventually occupying the entire cytosolic space (≥100 bacteria/cell). By contrast, vacuolar bacteria display low to moderate replication (2-40 bacteria/cell). Comparing the three treatment conditions, the incidence of infected cells with ≥100 bacteria at 8 h p.i. (8.2±1.6% in control, n=3 experiments) was reduced by EBSS treatment (1.2±2.0%) and promoted by WTM treatment (16±5.4%) (Fig 1D). Neither bacterial invasion at 1 h p.i. nor vacuolar replication at 8 h p.i. was overtly affected (Fig 1D). In agreement, total bacterial proliferation was restricted and promoted by EBSS and WTM treatment, respectively, compared to untreated cells (Fig 1E). Collectively, these data confirm that modulation of autophagy selectively affects the proliferation of *S*. Typhimurium in the cytosol of epithelial cells.

### Identification of cytosol- and vacuole-specific genes by RNA-seq analysis

To define the transcriptional signature of cytosolic *S*. Typhimurium, we isolated bacterial RNA from epithelial cells that had been enriched for either cytosolic or vacuolar S. Typhimurium by EBSS- or WTM-treatment, respectively, for transcriptomic analysis. Treated cells were infected with wild-type bacteria (Fig 1A), samples were collected and processed at 8 h p.i. and bacterial RNA was extracted using an enrichment protocol described previously ([31]; Fig S1). Four cDNA libraries were derived from two biological replicates and analyzed by RNA-seq (see Materials and Methods; [32]). The entire dataset is available in Dataset S1 and includes the expression profiles of the 4,742 coding genes and 280 sRNA identified previously in *S*. Typhimurium SL1344 [32, 33]. We defined genes/sRNAs as preferentially induced in the cytosol (“up cytosol”, WTM/EBSS ≥1.40-fold change) and vacuole (“up vacuole”, WTM/EBSS ≤0.61-fold change), respectively, and identified 113 statistically-significant “up cytosol” genes/sRNAs and 242 “up vacuole” genes/sRNAs (Dataset S1).

Of the twelve chromosomally-encoded pathogenicity islands (PAIs) in *S*. Typhimurium, only four were up-regulated in the cytosolic environment (Fig S2). Most obvious was the abundance of SPI1 genes that were up-regulated, consistent with the reported induction of *prgH* in the cytosolic population at later times in epithelial cells [12, 34]. SPI4, which encodes a giant non-fimbrial adhesin and its cognate type I secretion system and is co-regulated with SPI1 [35, 36], was also up-regulated in the cytosol. Lastly, genes encoding for two T3SS1 effectors within SPI5 and SPI11, SopB [37, 38] and SopF [22,23,39], respectively, were up-regulated in cytosolic bacteria. The remaining SPI5 and SPI11 genes were either up-regulated in the vacuolar population, or had unchanged expression, highlighting the mosaic nature of PAIs. Most SPI2 genes were up-regulated in the vacuolar population, as were genes encoding two type III effectors translocated by T3SS2 (*pipB* in SPI5 and *sspH2* in SPI12). SPI3 genes were up-regulated in the SCV of epithelial cells, like in macrophages [40, 41]. Similarly, most genes within SPI11, a PAI that is important for macrophage survival [42], and SPI12 were up-regulated in the epithelial SCV. The majority of SPI6 genes (which encode a type VI secretion system; [43]), all SPI9 genes (encode a type I secretion system; [44]) and SPI16 genes were not expressed in bacteria residing within either niche in epithelial cells. Overall, the transcriptional signatures of cytosolic and vacuolar bacteria at the PAI level (Fig S2) validates the basis of our RNA-seq analysis.

To put the observed gene expression changes into the context of the extensive *S. enterica* literature, we used an innovative approach that involved the curation of 80 custom pathways from previously published regulons that had been defined by RNA-seq, ChIP-seq or microarray-based analysis. We combined these pathways with KEGG and GO gene sets to annotate the subsets of “up cytosol” and “up vacuole” *S*. Typhimurium genes/sRNAs (Fig 2; Dataset S2). The list of “up vacuole” genes/sRNAs included an abundance (e.g. SPI2, *pipB*, *pgtE*, PinT, InvS, *steB*) that are positively regulated by the two-component systems that govern SPI2 induction in the SCV, namely PhoPQ, SsrAB and OmpRZ. Conversely, genes/sRNAs that are negatively regulated by these two component systems were in the “up cytosol” category. The SPI1-encoded transcription factors, HilA, HilD, and InvF, are required for the expression of SPI1 genes [45], and accordingly many of the “up cytosol” genes/sRNAs (e.g. *sipBCDA*, *siiCD*, *sopF*, *sopB*) grouped with these regulons. Our analysis also revealed many specific gene signatures (Fig 2). For example, genes associated with iron transport were solely up-regulated in the cytosol. Up-regulation of the tricarboxylic acid (TCA) cycle, lysine degradation and molybdate transport was evident in vacuolar bacteria, suggesting increased reliance of *Salmonella* on these energy sources in the SCV. Interestingly, genes/sRNAs that are repressed by Ferric uptake regulator (Fur) in iron-replete conditions and activated by Hfq, BarA/SirA and FliZ were up-regulated in both the vacuole and cytosol.

**Fig 2.**
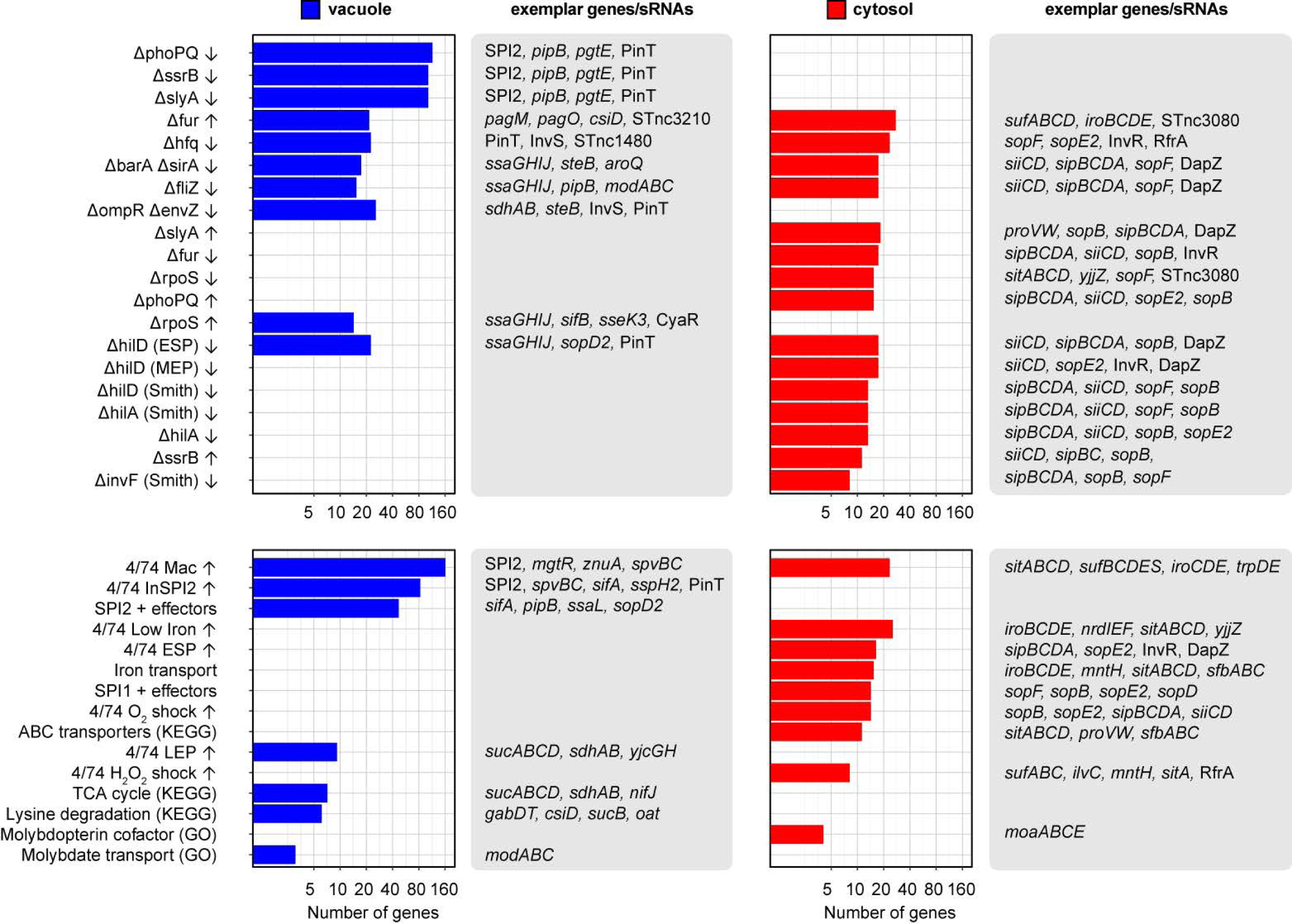
Pathway enrichment analysis results for the RNA-seq analysis. Top panels indicate regulons generated from transcriptional profiling of key regulatory mutants, while bottom panels display enriched gene sets generated from KEGG, GO, or from RNA-seq profiling of *S.* Typhimurium under 22 infection-relevant *in vitro* growth conditions. All displayed enrichments are significant (adjusted p-value <0.05) according to Fisher’s exact test with Benjamini-Hochberg correction for multiple comparisons. The number of genes/sRNAs in each category is shown in log_2_ scale. Only a selection of non-redundant pathways is presented; the full data set is available in S2 Table.

We used the expression profiles of *S*. Typhimurium grown in 22 distinct infection-relevant *in vitro* growth conditions [32] to predict the environmental conditions experienced by *S*. Typhimurium when in the cytosol and vacuole. For the 113 genes/sRNAs up-regulated in the cytosol, three main patterns emerged: (i) up-regulation by bile shock (3% bile for 10 min) and/or low Fe^2+^ shock (200 µM 2,2-dipyridyl (DPI) for 10 min) (e.g. *mntH*, *sitABCD*, *entCEBA*), (ii) up-regulation by NaCl shock (0.3 M NaCl for 10 min) (e.g. *proVW*, *ilvX*), (iii) up-regulation during late exponential/early stationary phase of growth (e.g. *sopE2*, *sipBCDA*, *siiABCDEF*) (Dataset S1). This suggests that cytosolic bacteria are growing fast and exposed to a low iron, high osmolarity physicochemical environment. The rapid growth of cytosolic *S*. Typhimurium has been previously observed [12,13,30], validating the latter prediction. A common theme observed for the 242 genes/sRNAs up-regulated in the vacuole was induction by: (i) *in vitro* SPI2 growth conditions and/or (ii) internalization into macrophages (Dataset S1).

### Cytosolic iron-limitation is a cue for *S*. Typhimurium gene/sRNA induction

To validate the cytosolic expression of genes/sRNAs predicted by RNA-seq analysis using *gfpmut3* transcriptional fusions because green fluorescent protein (GFP) is an ideal reporter to study differential gene expression in individual bacteria in infected host cells and tissues [12,17,46,47]. Previously *gfpmut3* transcriptional fusions have demonstrated the vacuole-specific induction of SPI2 genes [48], for example. We initially focused on those genes/sRNAs that are induced when *Salmonella* are exposed to low Fe^2+^ and/or bile shock in broth [32] (Dataset S1). Epithelial cells were infected with mCherry-*S*. Typhimurium carrying individual *gfpmut3* reporters and bacterial GFP fluorescence intensity was assessed qualitatively and quantitatively at 8 h p.i. The following genes/sRNAs were confirmed to be up-regulated in the cytosol: *entC* (*entCEBA* operon), *fepA* (*fepA-entD* operon), *fepB*, *fhuA* (*fhuACBD* operon), *fhuE*, *iroB* (*iroBCDE* operon), *iroN*, *mntH*, *nrdH* (*nrdHIEF* operon), *sitA* (*sitABCD* operon), SL1802, SL3990 (SL3990-SL3989 operon), *sufA* (*sufABCDSE* operon), *yjjZ* (SL1344_4483), STnc3080 and STnc3250 (Fig 3, upper panels; Fig S3). Up-regulation of STnc4000, *ilvC* and SL1344_2715 was not observed by fluorescence microscopy (Fig S6). Quantification using ImageJ software of the mean fluorescence intensity (MFI) of the GFP signal associated with individual bacteria confirmed the qualitative analysis for *iroB* (11.2-fold increase in MFI for cytosolic bacteria), *nrdH* (10.6-fold), *sitA* (13.3-fold), *sufA* (Fig 6.2-fold), *yjjZ* (23.2-fold) and STnc3080 (11.1-fold) (Fig 3, lower panel).

**Fig 3.**
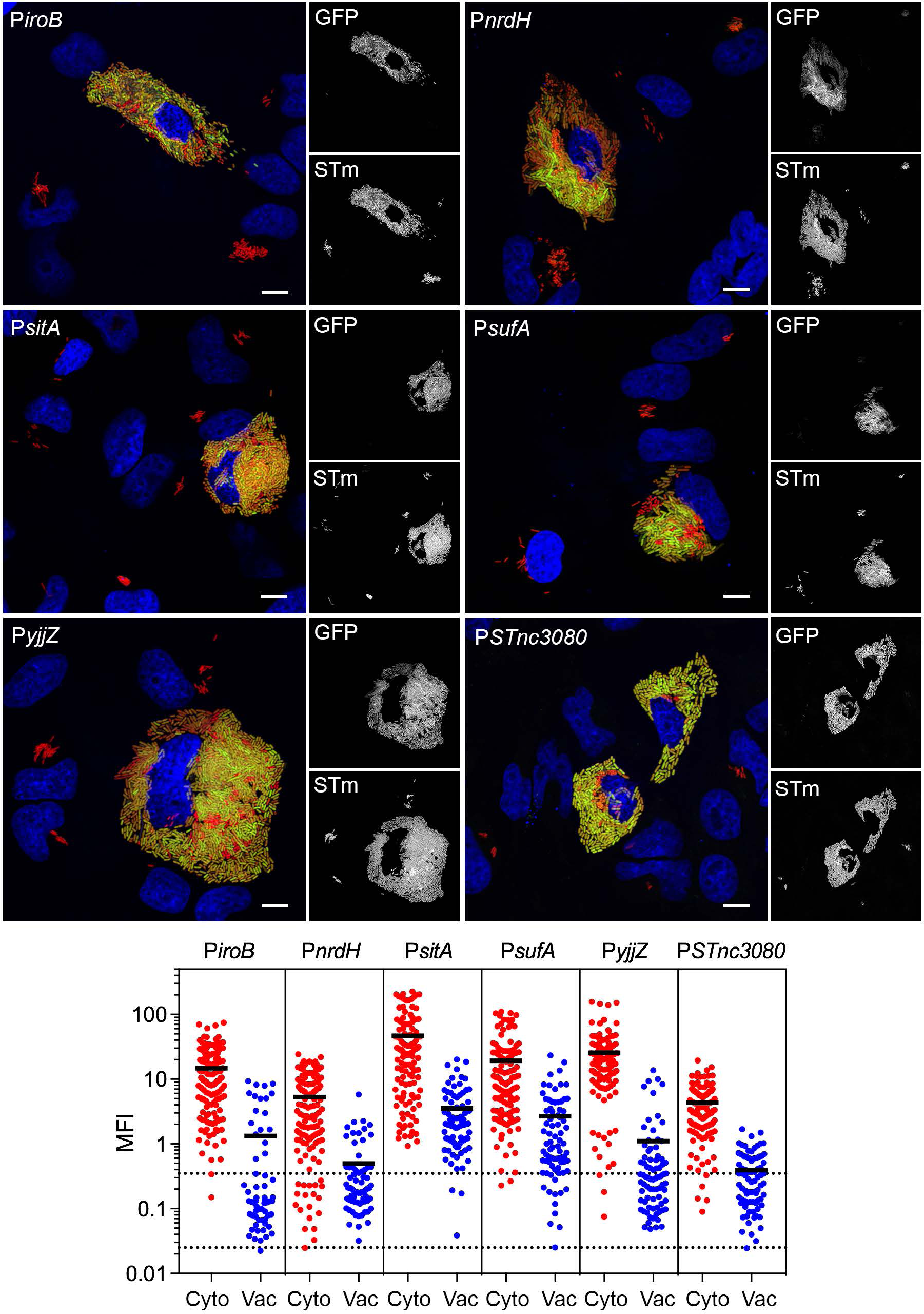
Iron-associated genes/sRNAs are induced in the epithelial cytosol. (A) Epithelial cells seeded on coverslips were infected wild-type mCherry-*S*. Typhimurium harboring fluorescent transcriptional reporters. The promoter region of each gene of interest was inserted upstream of a promoterless *gfpmut3*. At 8 h p.i., cells were fixed and stained with Hoechst 33342 to detect DNA. Representative confocal microscopy images show induction of *iroB*, *nrdH*, *sitA*, *sufA*, *yjjZ* and *STnc3080* promoters in cytosolic bacteria. Green = transcriptional reporter, red = *S*. Typhimurium, blue = DNA. Scale bars are 10 µm. (B) Quantification of the mean fluorescence intensity (MFI) of GFP signal in individual bacteria. Bacteria were designated as being cytosolic (Cyto) or vacuolar (Vac) if residing within cells with ≥100 bacteria or 2-40 bacteria, respectively. Each dot represents one bacterium. Solid black line indicates the mean. Acquisition parameters (acquisition time and gain) were set up using P*sitA-gfpmut3* (the highest GFP intensity) and these same parameters were applied throughout. The MFI for each bacterium was determine using ImageJ. Dashed lines indicate the range of background fluorescence in the GFP channel measured for mCherry-*S*. Typhimurium (no reporter). n=2 independent experiments.

All of the confirmed genes/sRNAs are up-regulated by “low Fe2+ shock” in broth (addition of 200 µM DPI for 10 min (35)). DPI chelation is not specific to iron, however [49, 50], raising the possibility that other metals also regulate the expression of these genes/sRNAs. We used *iroN*, *sitA*, *yjjZ*, *sufA*, *fepA* and STnc3250 promoters to determine metal cation specificity *in vitro*. The effect of increasing concentrations (0.1-100 µM) of Co^2+^, Fe^3+^, Mn^2+^, Ni^2+^ or Zn^2+^ on GFP fluorescence for bacteria grown in defined minimal media (M9) was measured (Fig S4). For all reporters, maximal expression was observed in the absence of metal addition. Upon addition of Fe^3+^ and Co^2+^, GFP expression of *S*. Typhimurium carrying P*iroN-gfpmut3*, P*yjjZ-gfpmut3* and P*fepA-gfpmut3* reporters decreased in a dose-dependent manner. Notably, the promoter responses were 10- to 100-fold more sensitive to Fe^3+^ than Co^2+^. P*sitA-gfpmut3* expression was modulated by Fe^3+^, Mn^2+^ and Co^2+^, although Fe^3+^ and Mn^2+^ were clearly the most potent repressors by an order of magnitude, in agreement with a previous study using a *sitA*::*lacZ* reporter [51]. Lastly, expression of P*STnc3250-gfpmut3* and P*sufA-gfpmut3* were only reduced by increased concentrations of Fe^3+^. P*iroN-gfpmut3* was the most sensitive iron reporter, with a >50% reduction in fluorescence at 0.1 µM Fe^3+^. Collectively, the *in vitro* repression of these genes/sRNAs by 0.1-1 µM Fe^3+^ hinted that the cue for cytosol-specific induction was iron limitation (<0.1 µM).

To test the response of reporters to iron concentrations during infection, we used ferric ammonium citrate (FAC) to increase free iron levels in mammalian cells [52, 53]. Epithelial cells were incubated overnight in growth media containing increasing concentrations (10-300 µM) of FAC. Untreated and FAC-treated cells were infected with mCherry-*S*. Typhimurium carrying transcriptional reporters, fixed at 8 h p.i. and the MFI of GFP signal for cytosolic and vacuolar bacteria quantified by fluorescence microscopy and ImageJ (Fig 4). The MFI of cytosolic bacteria harboring P*iroN-gfpmut3*, P*sitA-gfpmut3* or P*yjjZ-gfpmut3* reporters was reduced in a dose-dependent manner over the FAC concentration range. Specifically, fluorescence was reduced by 56.7-, 35.9- and 14.5-fold, respectively, upon the addition of 300 µM FAC to growth media (Fig 4A), indicating repression of *iroN*, *sitA* and *yjjZ* expression in the cytosol under iron-replete conditions. The low MFI for vacuolar bacteria in untreated cells limited our analysis of the iron-dependent repression of *iroN*, *sitA* and *yjjZ* in the SCV.

**Fig 4.**
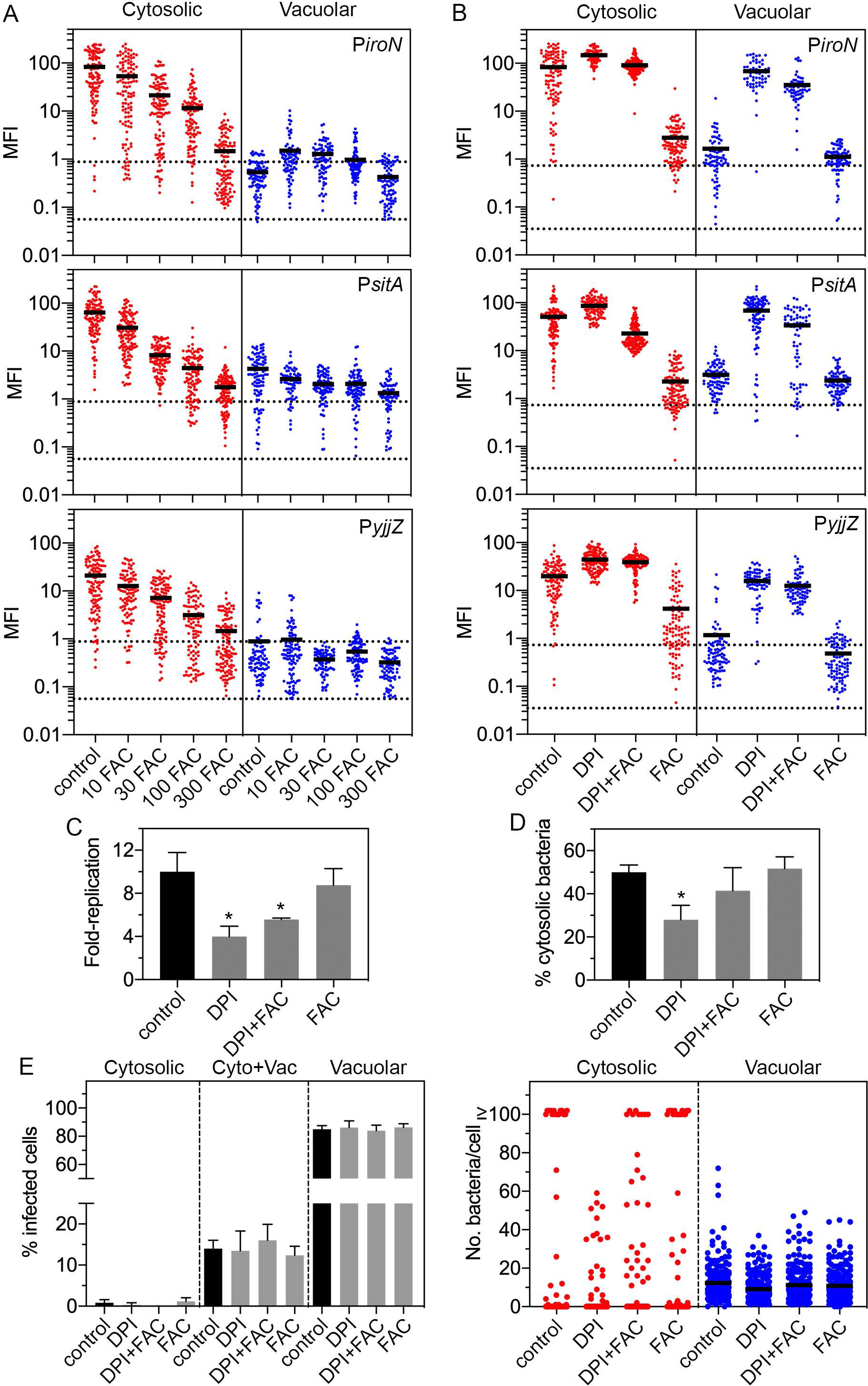
Manipulation of cellular iron levels primarily affects cytosolic bacteria. (A) Epithelial cells seeded on coverslips were left untreated (control) or treated overnight in growth media supplemented with 10 µM, 30 µM, 100 µM or 300 µM ferric ammonium citrate (FAC). Cells were infected with wild-type mCherry-*S*. Typhimurium harboring P*iroN-gfpmut3*, P*sitA-gfpmut3* or P*yjjZ-gfpmut3* reporters, fixed at 8 h p.i. and the MFI of GFP signal in individual bacteria was quantified by fluorescence microscopy. Bacteria were designated as being cytosolic (Cyto) or vacuolar (Vac) if residing within cells with ≥100 bacteria or 2-40 bacteria, respectively. Each dot represents one bacterium. Solid black line indicates the mean. Acquisition parameters (acquisition time and gain) were set-up using P*iroN-gfpmut3* (the highest GFP intensity) and these same parameters were applied throughout. The MFI for each bacterium was determine using ImageJ. Dashed lines indicate the range of background fluorescence in the GFP channel measured for mCherry-*S*. Typhimurium (no reporter). n=2 independent experiments. (B) Epithelial cells seeded on coverslips were left untreated (control) or treated overnight with growth media containing 200 µM 2,2-dipyridyl (DPI), a metal-chelating compound, 200 µM DPI + 200 µM ferric ammonium citrate (DPI+FAC) or 200 µM FAC alone (FAC). Infection strains and quantification of GFP signal were as described for (A). n=2 independent experiments. (C) Cells were treated as in (B) and infected with wild-type bacteria. The number of intracellular bacteria was quantified by gentamicin protection assay at 1 h and 8 h p.i. Fold-replication is CFUs at 8 h/1 h. n=3 experiments. (D) Cells were treated as in (B) and infected with wild-type bacteria. The proportion of cytosolic bacteria was quantified by CHQ resistance assay at Fig 6 h p.i. n≥3 experiments. (E) Cells were treated as in (B) and infected with mCherry-*S*. Typhimurium harboring a plasmid-borne reporter of cytosolic access, pNF101, and fixed at 8 h p.i. Left panel: the proportion of infected cells containing only cytosolic (all bacteria are GFP-positive, mCherry-positive), only vacuolar (all bacteria are GFP-negative, mCherry-positive) or a mixed population (Cyto+Vac) of bacteria was blindly scored by fluorescence microscopy. n≥5 independent experiments. Right panel: the number of cytosolic or vacuolar bacteria in each cell was scored. Horizontal black lines indicate the mean number of bacteria/cell. Cumulative data from two independent experiments is shown.

We used the lipophilic, cell-permeable chelator, DPI, as an independent way to assess the effects of metals on *S*. Typhimurium gene expression in the mammalian cytosol. Because DPI has a high affinity for various metal cations, we assessed the specificity for iron by means of an “add-back” experimental design [49,54,55] whereby an equimolar amount of Fe^3+^ was added to the DPI-treated media. Epithelial cells were treated overnight with 200 µM DPI, 200 µM DPI plus 200 µM FAC, or 200 µM FAC alone and infected with mCherry-*S*. Typhimurium harboring P*iroN-gfpmut3*, P*sitA-gfpmut3* or P*yjjZ-gfpmut3* reporters. Cells were fixed at 8 h p.i. and the intensity of GFP signal for cytosolic and vacuolar bacteria quantified via fluorescence microscopy. Addition of DPI increased the expression of *iroN*, *sitA* and *yjjZ* for both cytosolic and vacuolar bacteria (Fig 4B), indicating that DPI accesses both cellular compartments. However, the effects of divalent cation chelation were more pronounced for the vacuolar population (13- to 43-fold increase in MFI) than the cytosolic population (1.7- to 2.2-fold increase in MFI). Add-back of iron restored the MFI to that of untreated cells for cytosolic bacteria harboring P*iroN-gfpmut3* and P*sitA-gfpmut3* reporters (Fig 4B), demonstrating that iron limitation is the primary stimulus for the up-regulation of these genes in the cytosol. Add-back of iron had negligible effects on the MFI of bacteria harboring the P*yjjZ-gfpmut3* reporter, regardless of their intracellular niche (Fig 4B), possibly due to a lower sensitivity of *yjjZ* to changes in free iron levels (Fig 4A) or a responsiveness to deprivation of metals other than iron in the intracellular environment (Fig S4). Add-back of iron only partially restored the expression of *sitA* and *iroN* in vacuolar bacteria (Fig 4B) suggesting that *sitA* and *iroN* respond to multiple divalent cations in the SCV or FAC treatment does not significantly increase free iron levels in endocytic-derived compartments. Taken together, our data show that iron limitation in the mammalian cytosol is a major cue sensed by *S*. Typhimurium.

Using the same treatment conditions, we determined the role of iron limitation on bacterial replication in epithelial cells. As assessed by gentamicin protection assay (Fig 4C), total bacterial replication was limited by treatment with DPI (4.0-fold replication over 8 h) and DPI plus FAC (5.6-fold), but not FAC alone (8.8-fold), compared to untreated cells (10.0-fold). The chloroquine (CHQ) resistance assay, which quantifies the proportion of cytosolic bacteria in the total population, showed a similar profile (Fig 4D). Compared to untreated cells at Fig 6 h p.i (50±3.4% cytosolic bacteria), treatment with DPI reduced the levels of cytosolic bacteria to 28±6.Fig 6%. The proportion of cytosolic bacteria was increased to 42±11% upon add-back of iron (DPI+FAC), whereas it was indistinguishable from untreated cells for FAC alone (52±5.5% cytosolic).

It has been shown that epithelial cells contain either cytosolic bacteria, vacuolar bacteria or a mix of both populations at later times p.i. [30]. We used single-cell analysis to monitor the effect of metals on the distribution of these intracellular populations. Using mCherry-*S*. Typhimurium harboring pFN101 as a fluorescent reporter for cytosolic access, we analyzed the intracellular populations at 8 h p.i., in addition to the number of cytosolic (GFP+, mCherry+) and vacuolar (GFP-, mCherry+) bacteria within each infected cell. There was no significant difference in the frequency of epithelial cells that only contained cytosolic bacteria (0-1.2% of infected cells), only vacuolar bacteria (84.0-86.4% of infected cells) or a mixed population (12.4-16.0% of infected cells) between any of the treatments (Fig 4E, left panel), indicating that metal chelation does not alter the intracellular distribution of bacteria. However, the ability of *Salmonella* to replicate in the cytosol was noticeably affected (Fig 4E, right panel), in agreement with the CHQ resistance assay results. In untreated (control) cells, 6.4±0.89% of infected cells contained ≥100 cytosolic bacteria/cell. DPI treatment reduced proliferation of *S*. Typhimurium in the cytosol such that only 0.40±0.89% of infected cells contained ≥100 cytosolic bacteria/cell (p<0.05, n≥4 experiments). Add-back of iron (DPI+FAC) partially restored the hyper-proliferative capacity of *S*. Typhimurium in the cytosol (3.3±1.9%; p<0.05 compared to untreated, n≥4 experiments) whereas FAC treatment alone (7.8±1.7%) was no different than control. DPI also limited bacterial replication in the SCV (Fig 4E, right panel) as evidenced by a reduction in the mean number of vacuolar bacteria/cell at 8 h p.i. (14.0 for untreated and 10.4 for DPI-treated, p<0.05, n≥5 experiments). Add-back of iron restored vacuolar replication (12.5 bacteria/cell) whereas FAC alone (11.5 bacteria/cell) was not significantly different to untreated cells. In summary, iron restriction affects *S*. Typhimurium proliferation in both the cytosol and vacuole in epithelial cells but has a more profound effect in the cytosol.

### Zinc and magnesium are limiting in the vacuole but not the cytosol

We investigated whether metals other than iron affected intracellular *S*. Typhimurium gene expression in epithelial cells. *S*. Typhimurium encodes multiple transporters for divalent cations [56]. The up-regulation of *sitA* and *mntH* in cytosolic *S*. Typhimurium (Fig 3; Fig S3), which are induced by iron and manganese limitation (Fig S4) [51, 54] and encode high affinity Mn^2+^ transporters [57, 58], suggests that Mn^2+^ availability is low in the cytosol. Zn^2+^ transport in *S*. Typhimurium is dependent on a high affinity transport system, ZnuABC [59], and its accessory protein, ZinT [60, 61], and a low affinity ZIP family metal permease, ZupT [62, 63]. *zupT* is constitutively expressed [62], in agreement with our RNA-seq analysis (Dataset S1). By contrast, *znuA* and *zinT* are induced when zinc is limiting [64] and both genes are in the “up vacuole” shortlist (Dataset S1). We found that a P*zinT-gfpmut3* reporter was exquisitely sensitive to zinc concentrations, with a 54% and 99% reduction in GFP fluorescence observed upon the addition of 0.1 µM and 1 µM Zn^2+^ to minimal media, respectively (Fig S4). When epithelial cells were infected with mCherry-*S*. Typhimurium harboring P*zinT-gfpmut3*, GFP fluorescence was evident in vacuolar, but not cytosolic, bacteria at 8 h p.i. (Fig 5), indicating that low-zinc conditions prevail in the SCV (<0.1 µM). Quantification of the MFI of the GFP signal associated with individual bacteria confirmed the qualitative findings (5.8-fold increase in vacuole; Fig 5, lower panel).

**Fig 5.**
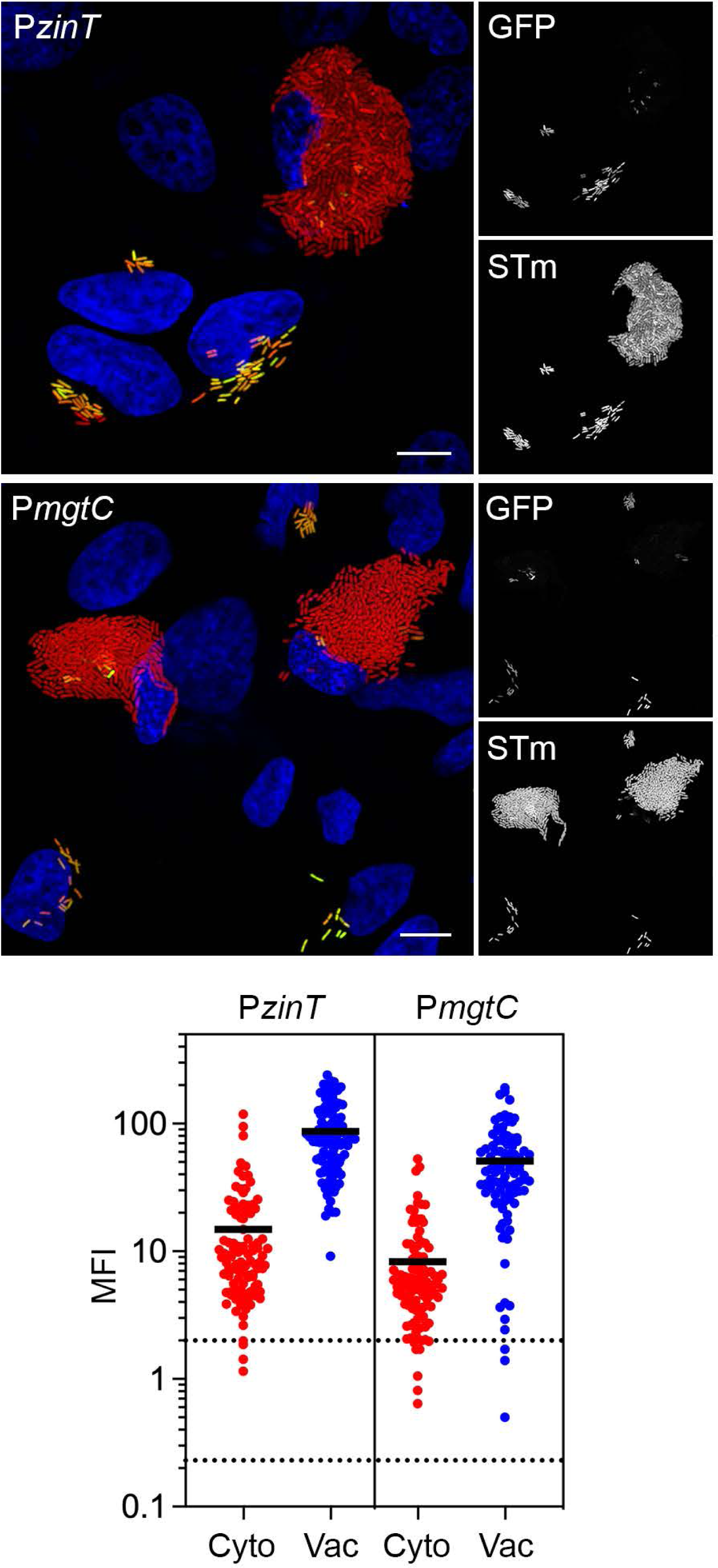
Zn^2+^ and Mg^2+^ are limiting in the vacuole, not the cytosol. Upper panels: Epithelial cells seeded on coverslips were infected with mCherry-*S*. Typhimurium harboring P*zinT-gfpmut3* or P*mgtC-gfpmut3* transcriptional reporters. At 8 h p.i., cells were fixed and stained with Hoechst 33342 to detect DNA. Representative confocal microscopy images show induction of *zinT* (upper panel) *and mgtC* (lower panel) promoters in vacuolar bacteria. Green = transcriptional reporter, red = *S*. Typhimurium, blue = DNA. Scale bars are 10 µm. Lower panels: Quantification of the MFI of GFP signal in individual bacteria. Bacteria were designated as being cytosolic (Cyto) or vacuolar (Vac) if residing within cells with ≥100 bacteria or 2-40 bacteria, respectively. Each dot represents one bacterium. Solid black line indicates the mean. Acquisition parameters (acquisition time and gain) were set-up using P*zinT-gfpmut3* (the highest GFP intensity) and these same parameters were applied throughout. The MFI for each bacterium was determine using ImageJ. Dashed lines indicate the range of background fluorescence in the GFP channel measured for mCherry-*S*. Typhimurium (no reporter). n=2 independent experiments.

**Fig 6.**
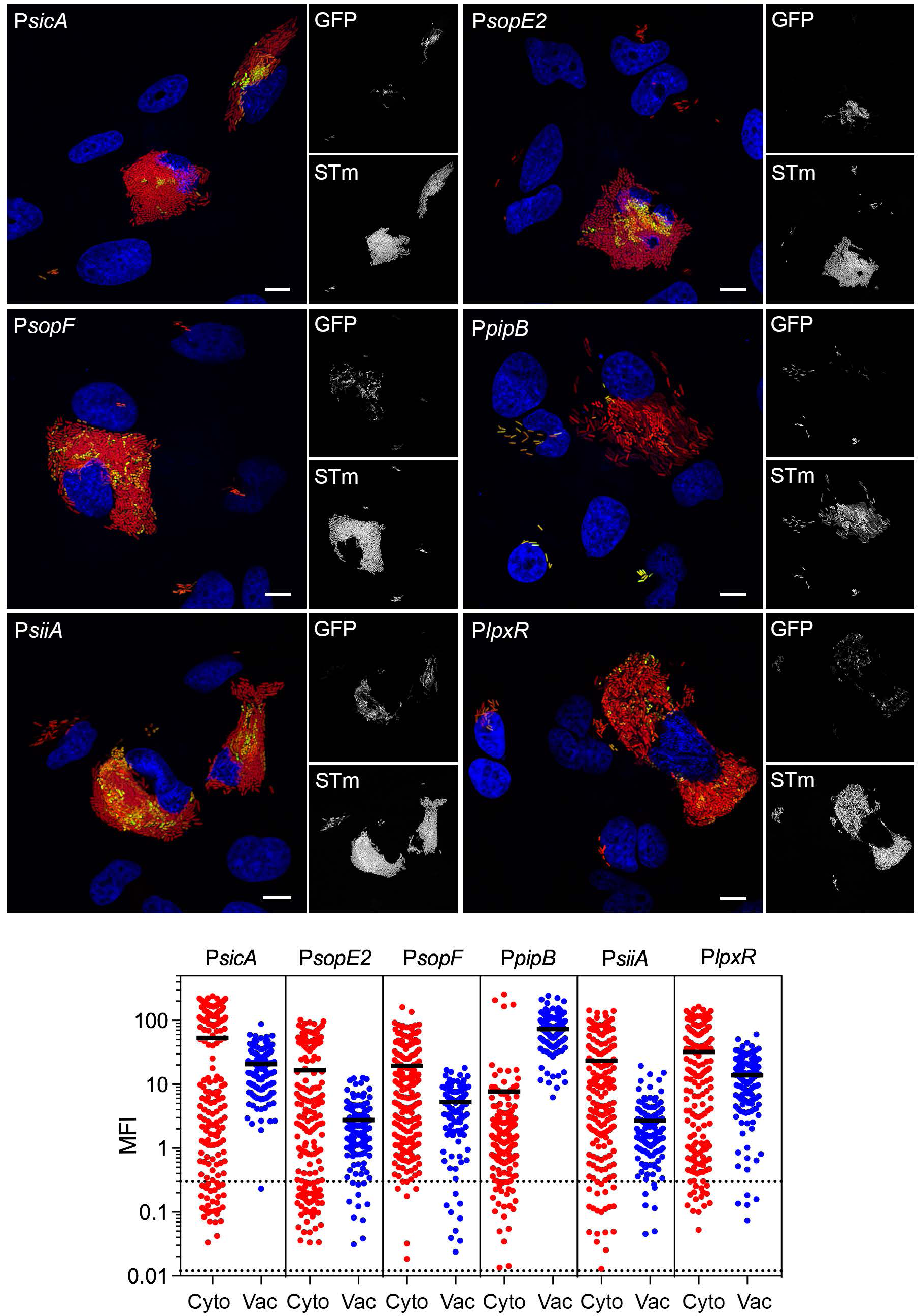
SPI1-associated genes are specifically up-regulated in the epithelial cytosol. (A) Epithelial cells seeded on coverslips were infected with mCherry-*S*. Typhimurium harboring *gfpmut3* transcriptional reporters. At 8 h p.i., cells were fixed and stained with Hoechst 33342 to detect DNA. Representative confocal microscopy images show induction of *sicA*, *sopE2*, *sopF*, *siiA* and *lpxR* promoters in cytosolic bacteria. PipB is a type III effector translocated by T3SS2 and the P*pipB-gfpmut3* reporter served as a control for vacuole-specific gene induction. Green = transcriptional reporter, red = *S*. Typhimurium, blue = DNA. Scale bars are 10 µm. (B) Quantification of the mean fluorescence intensity (MFI) of GFP signal in individual bacteria. Bacteria were designated as being cytosolic (Cyto) or vacuolar (Vac) if residing within cells with ≥100 bacteria or 2-40 bacteria, respectively. Each dot represents one bacterium; solid black line indicates the mean. Acquisition parameters (acquisition time and gain) were set-up using P*sicA-gfpmut3* (the highest GFP intensity) and these same parameters were applied throughout. The MFI for each bacterium was determine using ImageJ. Dashed lines indicate the range of background fluorescence in the GFP channel measured for mCherry-*S*. Typhimurium (no reporter). n=2 independent experiments.

*S.* Typhimurium also multiple Mg^2+^ transporters. CorA is a constitutively expressed Mg^2+^ transporter, and MgtA and MtgB are inducible Mg^2+^ transporters [65]. Consistent with its constitutive expression, expression of *corA* was similar in both vacuolar and cytosolic bacteria (Dataset S1). Low Mg^2+^ induces *mgtA* and *mgtB* transcription [65, 66]. Expression of *mgtA* was not induced in the vacuole, however (Dataset S1). Acidic pH abolishes the *mgtA* (but not *mgtB*) transcriptional response to Mg^2+^ concentration [66], which could explain why *mgtA* is not induced in the SCV of epithelial cells. In contrast, all three genes within the *mgtCBR* operon were up-regulated in vacuolar bacteria according to the RNA-seq data (Dataset S1). Using mCherry-*S*. Typhimurium harboring P*mgtC-gfpmut3*, we confirmed *mgtCBR* is responsive to Mg^2+^ levels (Fig S4) and the operon is specifically up-regulated in the SCV (Fig 5). The MFI of GFP signal was increased by 6.2-fold for vacuolar bacteria (Fig 5, lower panel). Altogether, our studies of Zn^2+^ and Mg^2+^ transporters reveal that levels of these cations are more limiting in the SCV than the cytosol.

### Confirmation of growth-phase related, cytosol-specific genes

Our transcriptomic analysis highlighted another set of cytosol-induced genes that were highly expressed during late exponential/early stationary phase of *in vitro* growth in broth, and in some cases responsive to oxygen shock (Fig 2; Dataset S1). Many of the genes encode for T3SS1 structural proteins (*orgA*, *spaQ*, *sipB*, *sipC*, *sipD*) and effectors known to be translocated by T3SS1 (*sipA*, *sopB*, *sopD*, *sopE*, *sopE2*, *sopF* [SL1344_1177]). Using GFP transcriptional reporters, we confirmed the cytosol-specific induction of *sicA* (*sicA-sipBCDA* operon), *sopE2* and *sopF* (Fig 6). We have previously shown that *prgH* (*prgHIJK-orgABC* operon), encoding a structural T3SS1 component, is up-regulated in cytosolic bacteria [12], in agreement with our RNA-seq results (Dataset S1). PipB, a T3SS2 effector encoded on SPI5 [67], was used as a control for the vacuole-specific induction of a type III effector. GFP fluorescence was only detected for vacuolar bacteria harboring a P*pipB-gfpmut3* reporter (Fig 6). We confirmed that *siiA* (*siiABCDEF* operon) and *lpxR* also showed cytosol-specific expression (Fig 6). The *siiABCDEF* operon in SPI4 (Fig S2) encodes for a type I secretion system for the secretion of SiiE, a giant non-fimbrial adhesin [68, 69]. Immunostaining with anti-SiiE antibodies detected the adhesin on the surface of cytosolic bacteria (Fig S5, upper panel), and diffusely spread throughout the cytosol (Fig S5, lower panel). Vacuolar bacteria were negative for SiiE staining (Fig S5). We have previously shown by electron microscopy that cytosolic bacteria have extensive filamentous material on their surface [12]. Taken together, our findings suggest the filaments could be SiiE. Like *siiABCDE*, *lpxR* (SL1344_1263) is also co-regulated with SPI1 [70] and encodes a hypothetical outer membrane protein. Transcriptional reporters did not verify cytosolic up-regulation of *ygbA* (SL1344_2840), which encodes a conserved hypothetical protein required for *S*. Typhimurium growth during nitrosative stress [71], or *asnA*, which is involved in asparagine biosynthesis [72] (Fig S6).

We conclude that there is a generalized up-regulation of “SPI1 associated” genes in the mammalian cytosol. However, we do not observe uniform up-regulation of these genes within the cytosolic population (Fig 6), in agreement with previous reports for *prgH* [12,14,34], which is different than that observed for the iron-responsive cytosol-specific genes/sRNAs (Fig 3). Such heterogeneity was verified by ImageJ quantification of the MFI of GFP signal associated with individual bacteria carrying P*sicA-gfpmut3*, P*sopE2-gfpmut3*, P*sopF-gfpmut3*, P*siiA-gfpmut3* or P*lpxR-gfpmut3* reporters. All genes were more highly expressed in cytosolic bacteria (Fig Fig 6, lower panel) – *sicA* (2.6-fold), *sopE2* (6.0-fold), *sopF* (3.7-fold), *siiA* (8.7-fold) and *lpxR* (2.3-fold) – yet with broader variability in MFI/bacterium than the iron-regulated genes (Fig 3), and with many cytosolic bacteria having background levels of GFP fluorescence.

### Extraneous genes that belong to the cytosol-specific program

Several osmotically-sensitive genes that are up-regulated when *S*. Typhimurium is exposed to NaCl shock [32] were identified as candidate cytosol-specific genes, namely *cysP* (*cysPUWA* operon), *soxS* and *proV* (*proVWX* operon) (Dataset S1). We qualitatively and quantitatively confirmed the cytosol-specific expression of *cysP* and *soxS* using transcriptional reporters (Fig S7). However, the *proV-gfpmut3* transcriptional fusion was not up-regulated in the cytosol (Fig S6). The *cysPUWA* operon encodes a sulfate/thiosulfate permease and periplasmic binding protein [73]. SoxS is a member of the AraC/XylS family of transcriptional regulators that is required for bacterial resistance to oxidative stress [74, 75].

Four additional *S*. Typhimurium genes were identified as being up-regulated in the cytosol according to the transcriptomics data, namely *uhpT*, *sfbA* (*sfbABC* operon), *mtr* and *trpE* (*trpEDCBA* operon) (Dataset S1). All four genes are constitutively expressed in infection-relevant broth conditions [32], and *sfbA*, *mtr* and *trpE* are also up-regulated upon infection of macrophages ([41]; Dataset S1). Infection of epithelial cells with mCherry-*S*. Typhimurium harboring P*uhpT-gfpmut3* or P*sfbA-gfpmut3* confirmed the RNA-seq-based prediction of their cytosol-specific induction (Fig S7). However, *mtr* and *trpE* were not induced in the cytosol (Fig S6). UhpT is a hexose phosphate transporter whose induction has been reported previously for cytosolic *S*. Typhimurium [34]. The *sfbABC* operon is predicted to encode a periplasmic iron-binding lipoprotein SfbA, a nucleotide-binding ATPase SfbB and a cytoplasmic permease SfbC [76].

### Identification of S. Typhimurium genes required for optimal cytosolic proliferation

Having identified *S*. Typhimurium genes/sRNAs that are specifically up-regulated within the epithelial cytosol, we next tested whether any of these genes/sRNAs are required for colonization of this niche. We did not focus on the SPI1-associated effectors as their individual contributions to the cytosolic stage of the intracellular cycle have been reported previously [22,30,77,78]. Eighteen gene deletion mutants were constructed and their ability to access and replicate within the cytosol was first assessed in a population-based assay by CHQ resistance. We identified five genes required for cytosolic replication in epithelial cells, namely *entC*, *fepB*, *sitA, mntH* and *soxS*. Deletion of these genes led to a reduced proportion of cytosolic bacteria at 7 h p.i. (42.5% for Δ*entC*, 41.9% for Δ*fepB*, 40.1% for Δ*mntH*Δ*sitA* and 42.6% for Δ*soxS*; Fig 7A). The remaining mutants had similar proportions of cytosolic bacteria as wild-type S. Typhimurium (50.2%, Fig 7A).

**Fig 7.**
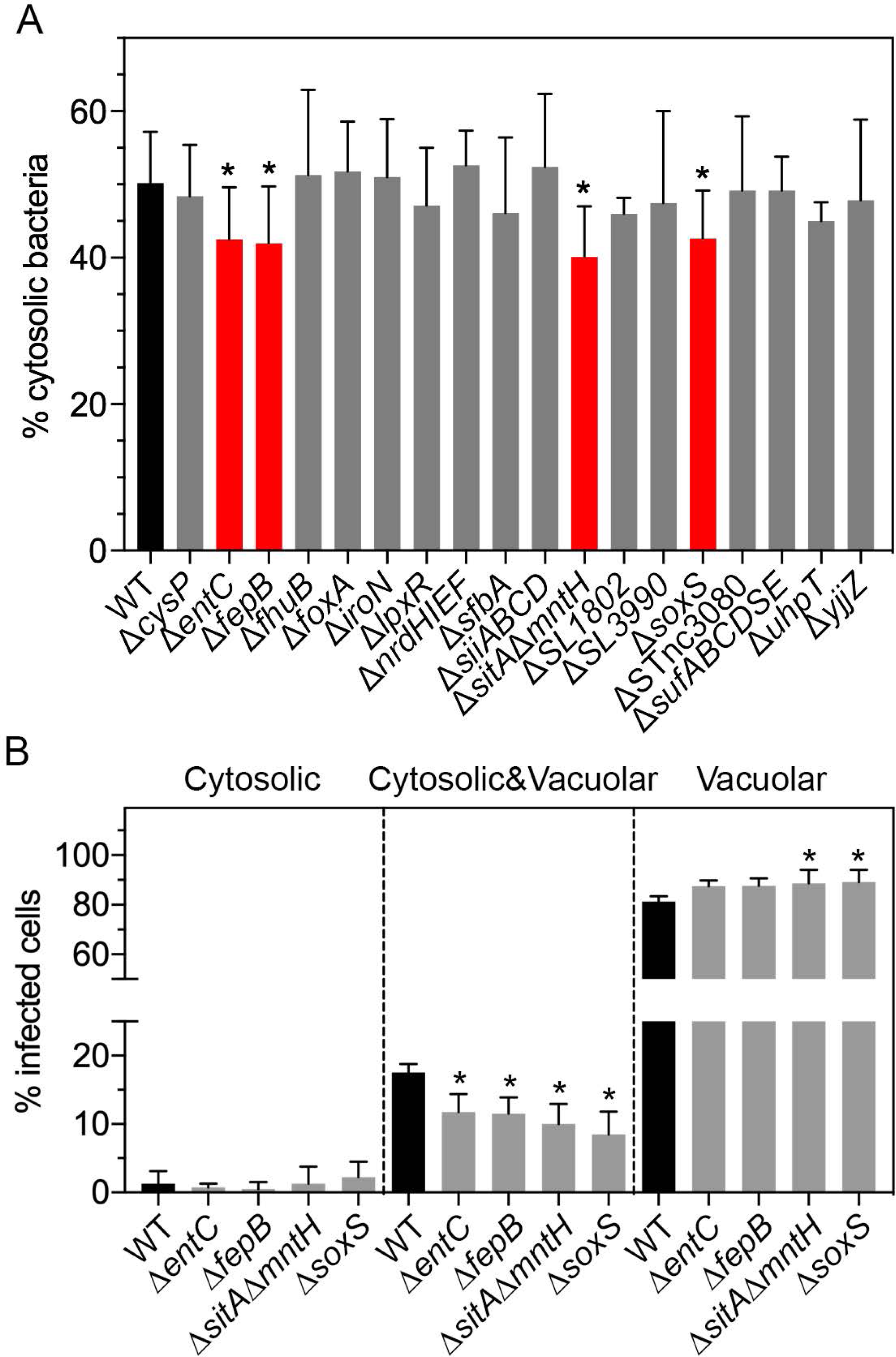
A subset of up-regulated genes is required for optimal proliferation in the cytosol. (A) Epithelial cells were infected with wild-type (WT) bacteria or the indicated gene deletion mutants and the proportion of cytosolic bacteria at Fig 6 h p.i. was quantified using the CHQ resistance assay. n≥5 independent experiments. (B) Epithelial cells seeded on coverslips were infected with WT bacteria or gene deletion mutant harboring pCHAR-Duo(ASV), a plasmid-borne dual reporter – constitutive *mCherry* expression is driven by the synthetic ProB promoter and *gfpmut3.1(ASV)* (encoding for destabilized GFP) is under the control of the glucose-6-phosphate responsive *uhpT* promoter from *S*. Typhimurium. The proportion of infected cells at 8 h p.i. containing only cytosolic (all bacteria are GFP-positive, mCherry-positive), only vacuolar (all bacteria are GFP-negative, mCherry-positive) or a mixed population (cytosolic and vacuolar) of bacteria was blindly scored by fluorescence microscopy. n=4 experiments.

We extended this population-based assay with single-cell analysis using *S*. Typhimurium harboring the dual fluorescence reporter plasmid, pCHAR-Duo(ASV), in which constitutive expression of *mCherry* is driven by the synthetic promoter ProB and destabilized GFP (*gfpmut3.1(ASV)*) is under the control of the *S*. Typhimurium *uhpT* promoter. All bacteria harboring this plasmid are mCherry-positive and only those bacteria in the cytosol will be GFP-positive. Epithelial cells were infected with wild-type, Δ*entC*, Δ*fepB*, Δ*sitA*Δ*mntH* or Δ*soxS* bacteria harboring pCHAR-Duo(ASV) and the number of infected cells containing only cytosolic bacteria, only vacuolar bacteria or a mixed population (cytosolic and vacuolar) at 8 h p.i. was scored by fluorescence microscopy. For all the deletion mutants, we detected fewer infected epithelial cells in the mixed population category than for wild-type bacteria (Fig 7B). Furthermore, fewer infected cells contained ≥100 cytosolic bacteria at 8 h p.i. (wild-type bacteria = 6.25%, Δ*entC* = 3.75%, Δ*fepB* = 5.0%, Δ*sitA*Δ*mntH* = 4.0% and Δ*soxS* = 4.0%; n=4 experiments). Altogether, these results identify *entC*, *fepB*, *sitA-mntH* and *soxS* as being specifically required for the optimal proliferation of *S*. Typhimurium in the cytosol of epithelial cells.

## Discussion

Compared to an endocytic-derived vacuole, the host cell cytosol could naïvely be viewed as a nutrient-rich milieu that allows for the efficient growth of bacteria. However, not all pathogens can survive within this niche, and *S*. Typhimurium can only replicate in the cytosol of epithelial cells and certain embryonic fibroblast lines [12,14,19]. Our findings that vastly different virulence gene programs are activated when *S*. Typhimurium colonizes the SCV and cytosol confirm previous suggestions that the mammalian cytosol is a complex environment that requires pathogen-specific adaptations for efficient bacterial survival and replication [79].

Here we report the “cytosol transcriptional signature” of bacteria by integrating our data with the available gene expression profiles of Gram-negative pathogens that colonize the cytosol (Table 1). We focus on the prototypical cytosolic pathogen, *S. flexneri* [80, 81], and uropathogenic *Escherichia coli* (UPEC) [82, 83], which forms intracellular bacterial communities (IBCs) in the cytosol of bladder epithelial cells. The most striking feature that emerges is the enrichment of genes that mediate the acquisition of iron (Table 1). *Salmonella* acquires ferric (Fe^3+^) iron by secreting two siderophores, enterobactin and salmochelin, a C-glucosylated form of enterobactin [56,84,85]. Iron-loaded siderophores then transit back into bacteria by first binding the outer membrane receptors, FepA and IroN [86, 87], then FepB in the periplasm, and finally across the cytoplasmic membrane via the ABC-type transporter, FepDGC [88]. FhuE, FhuA and FoxA also function as outer membrane receptors to exploit the utilization of ferric siderophores produced by other bacteria [89]. All of the siderophore synthesis and transport loci are up-regulated in cytosolic *S*. Typhimurium (Table 1, Fig 2). *S. flexneri* uses a similar strategy involving the synthesis of another siderophore, aerobactin [90], and the genes required for aerobactin synthesis and binding (*iucABCD* operon) are highly up-regulated in the cytosol (Table 1). The limited data available for UPEC IBCs are consistent with an important role of ferric iron acquisition for this bacterium (Table 1). Genes encoding the ferrous (Fe^2+^) iron transport systems of *S. flexneri* (*feoB*, *sitABCD* and *mntH*) and *S*. Typhimurium (*sitABCD* and *mntH*) are also up-regulated, but not *feoB* (Table 1). The Sit and MntH systems in *S*. Typhimurium transport both ferrous iron and manganese, but likely function as manganese transporters under physiological conditions [57, 58]. We found that a *S*. Typhimurium Δ*sitA*Δ*mntH* mutant is compromised for growth in the epithelial cytosol (Fig 7), indicating a specific requirement for Mn^2+^ acquisition in this compartment, but the same transport systems do not contribute to the growth of *S. flexneri* in epithelial cells [91]. The panopoly of iron acquisition systems used by enteric bacteria means that genetic deletions in multiple pathways are required to affect intracellular proliferation. For example, single gene deletions of *sitA*, *iucB* or *feoB* in *S. flexneri* have no effect on intracellular growth but a mutant in all three genes is unable to grow [92]. Similarly, *S*. Typhimurium Δ*entC* (defective in the synthesis of enterobactin and salmochelin; [93]) and Δ*fepB* (defective for the import of enterobactin, salmochelin and catecholate breakdown products containing iron; [84]) mutants fail to proliferate in the epithelial cytosol (Fig 7).

**Table 1.**
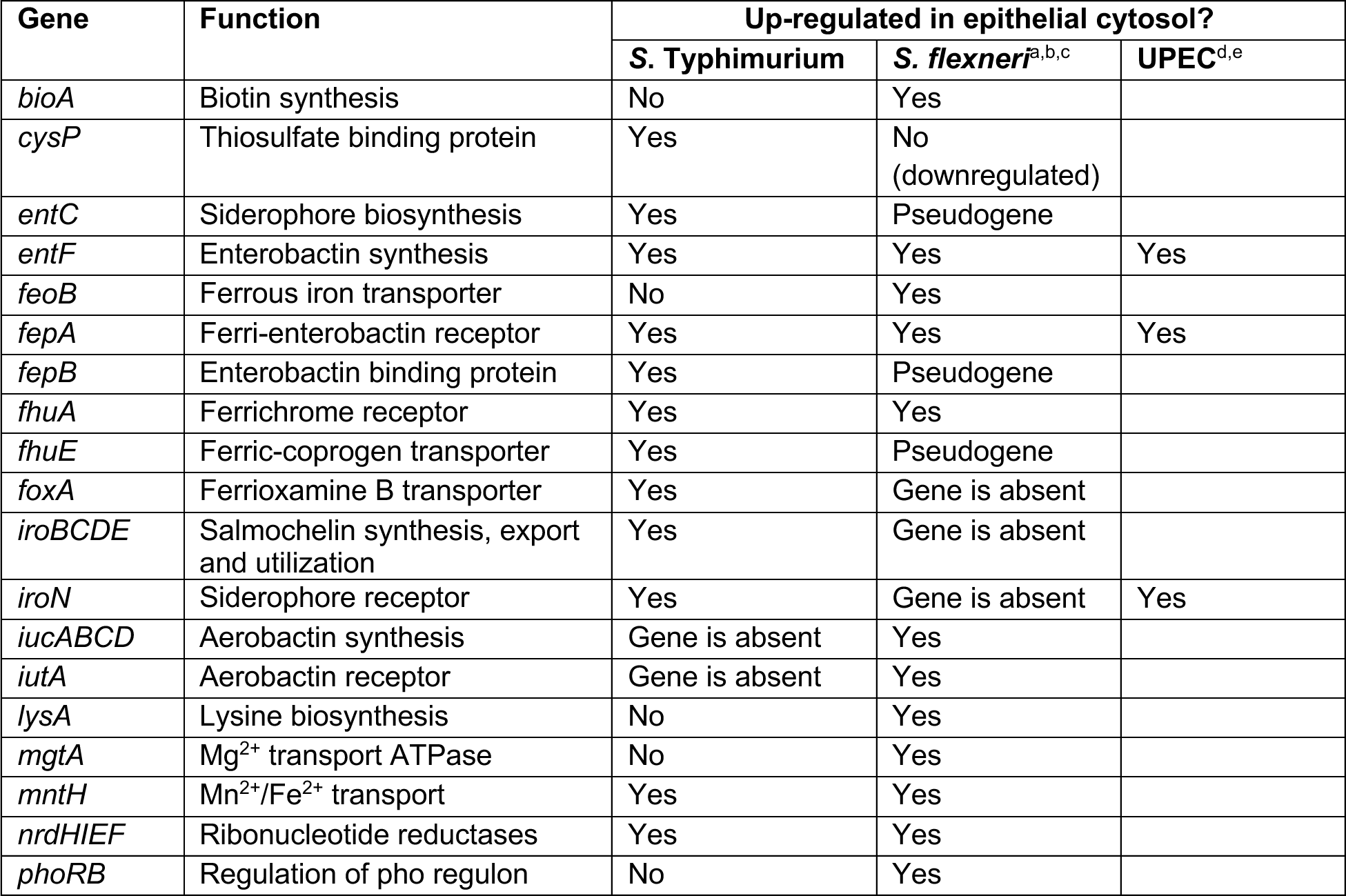
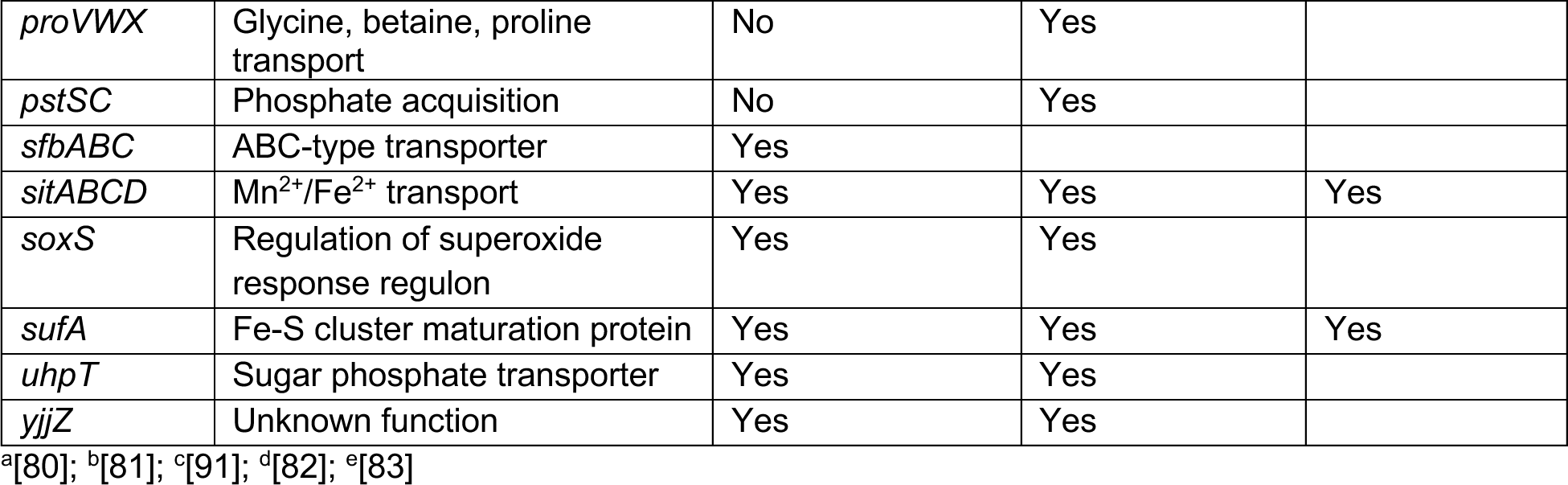
: Cytosol signature genes

We identified twelve other signature genes of cytosolic colonization: *nrdHIEF*, *sufABCDSE*, *yjjZ* and *uhpT* (Table 1). NrdEF is a Mn^2+^-dependent ribonucleotide reductase that functions under aerobic conditions, but only during iron restriction [94, 95]. The *suf* operon encodes for an iron-sulfur cluster system and is induced by oxidative stress and iron starvation [96], like the *nrdHIEF* operon [97, 98]. Fur is the master regulator of iron homeostasis in bacteria [99]. While the function of YjjZ remains unknown, *yjjZ* (SL1344_4483) is part of the Fur regulon in *S*. Typhimurium [100] and UPEC [101]. In summary, transcription of *nrdHIEF*, *sufABCDSE* and *yjjZ* is controlled by iron deprivation and Fur. The sole metabolic gene identified as part of the cytosol transcriptional signature is *uhpT*, a gene that is not induced by iron deprivation or oxidative stress but rather by glucose-6-phosphate levels in the cytosol [80].

The combination of available transcriptome data revealed bacterial species-specific cytosol-induced genes. Specifically, *bioA*, *lysA*, *pstSC*, *phoRB*, *proVWX* and *mgtA* were only induced in *Shigella* and not in *S*. Typhimurium (Table 1; Dataset S1; [81, 92]). The differential expression profile suggests variance in the metabolic and phosphate acquisition strategies used by these two pathogens in the epithelial cytosol. The contribution of Mg^2+^ transport also seems to differ between the two bacteria, as *mgtA*-encoded Mg^2+^ acquisition is only induced by the cytosolic environment for *S. flexneri*, and not for *S*. Typhimurium. Unlike *S. flexneri*, *S*. Typhimurium has a second Mg^2+^ transport system encoded by *mgtB*, which is only up-regulated in the vacuole (Dataset S1; Fig 5). *S. flexneri* does not encode for *mgtB*. Furthermore, the third Mg^2+^/Ni^2+^/Co^2+^ transporter (CorA) is constitutively expressed in *S*. Typhimurium [102, 103] and is required for optimal proliferation in the cytosol [104]. The importance of Mn^2+^ acquisition also seems to differ (see above).

We identified *S*. Typhimurium-specific genes/sRNAs that serve as signatures of the cytosolic lifestyle. Examples include STnc3080, STnc3250, SL1802 and *siiABCDEF*, as well as eight type III effectors that are translocated by T3SS1 (Fig 3, Fig 6, Figure S3, Dataset S1). STnc3080, STnc3250 and SL1802 are all negatively regulated by Fur [32,100,105] and have not previously been reported to play a role in *Salmonella* pathogenesis. Intracellular expression of the *siiABCDEF* operon has not been reported before, prompting us to establish that the SiiE protein, which mediates adhesion to the MUC1 on the apical surface of intestinal epithelial cells [106], is both on the surface of cytosolic bacteria and secreted into the cytosol of epithelial cells (Fig S5). The combination of findings from this study, and others [12, 34], shows that cytosolic *S*. Typhimurium are induced for T3SS1, secrete T3SS1-associated effectors, are flagellated and decorated with SiiE. We conclude that cytosolic *S*. Typhimurium are “primed” to enter naïve enterocytes following cellular release.

The major environmental factor that controls the transcriptional program of cytosol-residing bacteria is clearly iron-limitation. Within mammalian cells, the “labile iron pool” refers to the pool of chelatable and redox-active iron [107, 108]. In resting cells, the labile iron pool is estimated to be ∼1 µM, representing only a minor fraction of the total cellular iron (50-100 µM) [107, 108]. At a subcellular level, measurements have shown that the concentration of labile iron is lower in the cytosol than the nucleus, mitochondria or endosomes/lysosomes [109, 110], in agreement with our findings. We believe that consumption of free iron from the cytosol by rapidly growing *S*. Typhimurium further depletes the pool compared to uninfected/resting cells.

The uniform induction of iron-responsive *S*. Typhimurium genes/sRNAs in the cytosol (Fig 3, Figure S3) indicates a generalized bacterial response to iron deprivation. Fur is the major iron-responsive transcriptional regulator that controls iron homeostasis and oxidative stress defences in bacteria [99]. When bound to iron (i.e. in iron-replete conditions), Fur acts as both a repressor and activator of gene expression, either directly or indirectly, and modulates expression of ∼10% of the *S*. Typhimurium genome [100, 111]. Except for SL3990, all the low Fe^2+^/bile shock-regulated “up cytosol” genes/sRNAs (Dataset S1) that were validated using transcriptional reporters (Fig 3, Fig S3) are negatively regulated by Fur [89,100,112](Fig 2). Many have a Fur box in their promoter regions, the regulatory site to which iron-bound Fur binds, suggesting direct regulation. However, an unanswered question is why all Fur-repressed *S*. Typhimurium genes/sRNAs are not induced in the cytosolic population ([100]; Dataset S1). It is also unclear why SPI1-associated genes are induced in a subset of cytosol-exposed bacteria in an iron-limited environment, when expression of the SPI1 regulon is typically activated by iron-bound Fur under iron-replete conditions via HilD and H-NS [113, 114]. The regulation of SPI1 and its associated genes is complex [45], with numerous environmental signals feeding into the regulatory network including osmolarity, oxygen and bacterial growth phase, as well as iron. Bacterial cell-to-cell variation in the integration of all the relevant signals could explain the phenotypic heterogeneity observed for SPI1-associated gene expression in cytosolic bacteria. Such heterogeneity in T3SS1 gene expression has been reported previously in broth cultures [115, 116].

We have identified the site-specific cues encountered by *S*. Typhimurium in the distinct milieus of a vacuole versus the cytosol. *S*. Typhimurium residing within the epithelial cytosol have limited access to iron and manganese, are experiencing oxidative stress and utilize hexose-6-phosphate sugars as a carbon source. Other studies of *S. flexneri* and *L. monocytogenes* colonization suggested that the mammalian cytosol is a nutrient-rich environment of neutral pH with low concentrations of Na^2+^, Mg^2+^, Ca^2+^ and PO_4_^3-^ ions, but a high concentration of K^+^ [79,81,117,118]. In contrast, the mature SCV is acidic [7,119,120], has high levels of K^+^, Fe^2+^ and oxygen, but is limiting in Mg^2+^, PO_4_^3-^ and Fe^3+^, and *S*. Typhimurium are experiencing oxidative stress but not amino acid starvation [40,41,81].

Based on single-cell analysis, we can refine previous conclusions drawn from population-based studies of *S*. Typhimurium and epithelial cells. For example, in the landmark study of Hautefort *et al*. (2008) [31], the simultaneous induction of SPI1 and flagella within epithelial cells was reported, which at the time was puzzling. Single-cell microscopic analysis has solved this mystery by showing that the cytosolic population is solely responsible for the intracellular expression of both SPI1 and flagella ([12,14,34]; this study). Based on the intracellular induction of *iroA*- and *mgtB*-*lacZ* reporter fusions in MDCK (epithelial) cells, concentrations of Fe^2+^ and Mg^2+^ were thought to be low in the mature vacuole [121]. Additionally, a recent proteomic analysis of *S*. Typhimurium isolated from infected epithelial cells at 6 h p.i. suggested that bacteria are starved for numerous metals, including iron, manganese, and zinc [122]. Our findings show that metal limitation experienced by intracellular bacteria is niche-dependent; levels of iron and manganese are low in the cytosol, whereas zinc and magnesium are limiting in the vacuole.

Here, we demonstrate the importance of verifying results derived from population-based studies at the single-cell level. We suggest that the many studies that have identified individual *S*. Typhimurium or mammalian genes that influence bacterial proliferation in epithelial cells should be revisited, especially if the distinct replication niches were not considered. As heterogeneity in terms of bacterial proliferation and intracellular colonization site clearly influences the outcome of infection [123, 124], it is crucial that the study of host-pathogen interactions incorporates the investigation of individual bacteria, as well as host cells.

## Materials and Methods

### Bacterial strains and plasmids

*S*. Typhimurium SL1344 was the wild-type strain used in this study [125]. Gene deletion mutants were constructed using *sacB* allelic exchange, λ red recombinase technology or P22 transduction. SL1344 Δ*sitA* (deleted for amino acids 4-298), Δ*mntH* (deleted for amino acids 1-411), Δ*yjjZ* (ΔSL1344_4483, deleted for amino acids 4-Fig 66) and ΔSL1344_1802 (deleted for amino acids 4-53) deletion mutants were made via *sacB* negative selection. Two fragments of ∼1 kb upstream and downstream of the gene of interest were amplified from *S*. Typhimurium SL1344 genomic DNA using Phusion High-Fidelity DNA polymerase (Thermo Scientific) (primer sequences are listed in Table S1). The two fragments were combined for a second round of amplification by overlap extension PCR. The amplicon was then digested with appropriate restriction enzymes, ligated into the suicide vector pRE112 (Cm^R^) [126], and electroporated into *E. coli* SY327λpir. After sequence confirmation, the pRE112 plasmids were transferred to *E. coli* SM10λpir (Kan^R^) for conjugation into SL1344 wild-type or Δ*mntH* bacteria (for the Δ*sitA*Δ*mntH* double mutant). For the second recombination event, *sacB*-based counterselection on LB agar containing 5% (w/v) sucrose was used, and streptomycin-resistant, chloramphenicol-sensitive colonies were screened by PCR with primers outside the recombination region to confirm the deletion of each gene. *S*. Typhimurium SL1344 Δ*sufABCDSE*::kan, Δ*nrdHIEF*::kan, Δ*fhuB*::kan and Δ*fepB*::kan strains were generated using λ red recombineering technology [127]. λ red cassettes were amplified using pKD4 as a template with the oligonucleotide pairs listed in Table S1 and electroporated into *S*. Typhimurium SL1344 wild-type containing the temperature-sensitive helper plasmid, pKD46. Transformants were selected on LB agar plates containing streptomycin (100 µg/ml) and kanamycin (50 µg/ml). Verification of λ red promoted gene replacement was by PCR using a target-flanked primer and a kanamycin resistance gene-specific primer. Each mutation was transferred into a clean SL1344 wild-type background using P22 phage transduction. The following insertional mutants were constructed by P22 transduction into a SL1344 wild-type background from phage lysates prepared from the following *S*. Typhimurium mutants: Δ*entC*::kan (Raffatellu, unpublished), Δ*iroN*::kan [84], ΔSTnc3080::kan (Kroeger, unpublished), ΔSL1344_3990::kan [128], Δ*cysP*::kan [128], Δ*foxA*::kan [128], Δ*soxS*::kan [128], Δ*sfbA*::kan [128], Δ*uhpT*::kan [128], Δ*lpxR*::kan [128], Δ*siiABCD*::kan [129]. Selection was on LB agar plates containing kanamycin (50 µg/ml).

SL1344 wild-type bacteria constitutively expressing *mCherry* from the chromosome (*glmS*::*Ptrc-mCherryST*::FRT, denoted as mCherry-*S*. Typhimurium) or a plasmid (pFPV-mCherry) have been described previously [7, 130]. *mCherry* expression is driven by the constitutive *trc* promoter or *S*. Typhimurium *rpsM* promoter, respectively. The *PuhpT-gfpova* plasmid, pNF101 [22], was used as a biosensor for bacterial exposure to the mammalian cytosol. Expression of the unstable GFP variant, *gfp_ova* (protein half-life <60 min, [131]), is under the control of the glucose-6-phosphate responsive *uhpT* promoter from *S. flexneri*. Alternatively, bacterial access to the cytosol was assessed using the bidirectional reporter plasmid, pCHAR-Duo(ASV), whereby constitutive *mCherry* expression is driven by the synthetic ProB promoter and *gfpmut3.1(ASV)* expression is under the control of the *S*. Typhimurium *uhpT* promoter. To create pCHAR-Duo(ASV), the coding sequence for the stable GFP variant, *gfpmut3.1* (protein half-life >24 h), was excised from pCHAR-Duo [30] by XmaI/ApaI digestion and replaced with the coding sequence for a destabilized GFP variant, *gfpmut3.1(ASV)* (protein half-life ∼110 min), which had been released from pGFP(ASV) (Clontech) by XmaI/ApaI digestion.

To construct fluorescent transcriptional reporters, we extracted the precise transcriptional start sites from the online SalComMac database (http://tinyurl.com/SalComMac; [32, 41]). Approximately 500 bp of sequence upstream of the ATG start codon of each gene, or transcript start site of each sRNA, was amplified from SL1344 genomic DNA with the oligonucleotide pairs listed in Table S2. Amplicons were digested with XbaI/SmaI and ligated into the corresponding restriction sites of pGFPmut3.1 (Clontech). XbaI/ApaI digestion released the gene/sRNA promoter-*gfpmut3* fragment, which was then ligated into the corresponding sites of the low copy number plasmid, pMPMA3ΔPlac [132]. The *pipB* and *sopB* promoters were excised from pMPMA3ΔPlac-P*pipB*-GFP(LVA) and pMPMA3ΔPlac-P*sopB*-GFP(LVA), respectively [132], by digestion with XbaI/KpnI and ligated into the corresponding sites of pGFPmut3.1 and then transferred to pMPMA3ΔPlac as described above. The pMPMA3ΔPlac-P*uhpT*-*gfpmut3* construct has been described previously [133]. GFP reporter plasmids were electroporated into wild-type mCherry-*S*. Typhimurium.

### Reporter assays for metal sensitivity

For metal repression studies, wild-type mCherry-*S*. Typhimurium bacteria harboring *gfpmut3* fusions were grown shaking overnight (220 rpm) at 37°C in LB-Miller broth containing 100 µg/ml streptomycin and 50 µg/ml carbenicillin, then washed twice in M9 salts pH 7.0 (6 g Na_2_HPO_4_, 3 g KH_2_PO_4_, 0.5 g NaCl, 1 g NH_4_Cl per liter) supplemented with 1 mM MgSO_4_, 0.4% (w/v) glucose and 0.01% (w/v) histidine – hereafter referred to as M9-supplemented media – and resuspended in an equal volume of M9-supplemented media. Washed bacteria were diluted 1:200 in M9-supplemented media containing 100 µg/ml streptomycin and 50 µg/ml carbenicillin and the desired concentration of cobalt (II) chloride (Alfa Aesar, Puratronic®, 99.998%), iron (III) chloride (Acros Organics, 99+%), manganese (II) chloride (Alfa Aesar, Puratronic®, 99.999%), nickel (II) chloride (Alfa Aesar, Puratronic®, 99.9995%) or zinc chloride (Alfa Aesar, Puratronic®, 99.999%). All solutions were prepared fresh in MilliQ water. Bacteria were grown overnight at 37°C, shaking at 220 rpm for ∼16 h. GFP fluorescence was measured in black 96-well plates (Costar) using a TECAN SPARK plate reader (excitation wavelength of 485 nm, bandwidth 20 nm; emission wavelength of 535 nm, bandwidth of 20 nm). All readings were normalized for bacterial growth (OD_600_). The background fluorescence of mCherry-*S*. Typhimurium bacteria (no reporter plasmid) was subtracted from all readings.

### Mammalian cell culture

HeLa human cervical adenocarcinoma epithelial cells were purchased from American Type Culture Collection (ATCC, CCL-2) and used within 15 passages of receipt. Cells were maintained at 37°C and 5% CO_2_ in the growth medium (GM) recommended by ATCC i.e. Eagle’s minimum essential medium (EMEM, Corning) containing sodium pyruvate, L-glutamine and 10% (v/v) heat-inactivated fetal calf serum (FCS; Gemini Bio Products). Tissue culture plasticware was purchased from Thermo Scientific Nunc.

### Bacterial infection of mammalian cells

HeLa cells were seeded 24 h prior to infection at the following densities: (i) 5×10^4^ cells/well in 24-well tissue culture plates, (ii) 6×10^4^ cells/well on acid-washed glass coverslips in 24-well plates, or (ii) 2.6×10^6^ cells/15 cm tissue culture dish. T3SS1-induced bacterial subcultures were prepared in LB-Miller broth (Difco) [134] and cells were infected for 10 min with bacterial subcultures (MOI ∼50) as described [134]. To induce autophagy, HeLa cells were shifted to Earle’s Balanced Salt Solution (EBSS, Sigma) 3 h prior to infection and maintained in EBSS until 90 min post-infection. Thereafter, infected cells were switched back to regular growth medium. To inhibit autophagy, cells were incubated in growth media containing 100 nM wortmannin (WTM, Calbiochem) for 45 min prior to infection, continuing to 90 min post-infection, whereupon cells were transferred back to regular growth media. For manipulation of cellular cation levels, epithelial cells were treated 16-18 h prior to infection in growth media containing 200 µM 2,2’-dipyridyl (DPI, Sigma-Aldrich), 200 µM DPI and 200 µM ammonium iron (III) citrate (FAC, Alfa Aesar) or 200 µM FAC alone. DPI +/− FAC treatment continued throughout the infection. DPI was prepared as a 100 mM stock in ethanol and stored at −80°C for a maximum of 2 weeks. FAC was prepared extemporaneously in MilliQ water. Gentamicin protection and CHQ resistance assays (chloroquine diphosphate salt, Sigma) were as described previously [134]. Monolayers were solubilized with 0.2% (w/v) sodium deoxycholate (Sigma) and serial dilutions were plated on LB-Miller agar (Remel) for enumeration of colony forming units (CFUs).

### Bacterial enrichment and RNA extraction from infected cells

*S*. Typhimurium was enriched from infected HeLa cells (12 x 15 cm dishes for EBSS treatment, 6 x 15 cm dishes for WTM treatment) as described previously by Hautefort *et al*. (2008) [31] with minor modifications. Briefly, infected HeLa cells were washed once with 30 ml chilled phosphate-buffered saline (PBS), then lysed on ice for 10-20 min in 0.1% SDS, 1% acidic phenol, 19% ethanol in water (15 ml per 15 cm dish). Pooled lysates were collected into 50 ml conical tubes and centrifuged at 3,220 xg for 30 min, 4°C. After three washes in ice-cold wash buffer (0.1% phenol, 19% ethanol in water) (30 min, 3,220 xg, 4°C each wash), the remaining bacterial pellet was resuspended in <1 ml of wash buffer, transferred to an RNase-free microcentrifuge tube and centrifuged at 16,000 xg for 2 min. The supernatant was discarded, and the pellet resuspended in 1 ml TRIzol^®^ reagent (Life Technologies) by gently pipetting up-and-down (∼60 times), then transferred to −80°C for storage. Total RNA was extracted as described [32] and the RNA concentration quantified using a Nanodrop 2000 spectrophotometer (Thermo Scientific). RNA quality was analyzed using an Agilent Bioanalyzer 2100. RNA samples isolated from two independent experiments were sent to Vertis Biotechnologie AG (https://www.vertis-biotech.com) for RNA sequencing.

### RNA-seq data analysis

A total of 42 to 57 million sequence reads were generated from each sample on the Illumina HiSeq platform. For the RankProduct analysis, segemehl [135] was used to map against the reference genome sequence, resulting in 18.2 million uniquely mapped reads per sample on average. For the BitSeq analysis, mapping against the transcriptome was required. This was accomplished through Bowtie2 [136] and generated 7.1 million uniquely mapped reads per sample on average. Gene/sRNA expression values were calculated by the Transcripts Per Million (TPM) approach [137]. All genes/sRNAs with mapped reads were reported as TPM and a threshold TPM value of 10 was used as a cut-off to distinguish gene expression from background noise [32]. Genes/sRNAs were deemed differentially expressed using fold-change criteria of ≥1.40 (cytosol up-regulated) and ≤0.71 (vacuole up-regulated) for WTM/EBSS treatment conditions (cells highlighted in yellow in Dataset S1). Statistical significance was assessed using two independent platforms, Bayesian interference of transcripts from sequencing data (BitSeq; [138]) and rank products (RankProduct; [139, 140]). For Bitseq analysis, we arbitrarily set a Probability of Positive Log Ratio (PPLR) of ≥0.90 for “up cytosol” genes/sRNAs and ≤0.1 for “up vacuole” genes/sRNAs for statistical significance (cells highlighted in green in Dataset S1). For RankProduct analysis, the cutoff for differentially expressed genes was arbitrarily set to a percentage of false positive (pfp) value of ≤0.2 (cells highlighted in green in Dataset S1) for statistical significance.

### Pathway enrichment analysis

A collection of 80 custom pathways was generated from previous RNA-seq, ChIP-seq, and microarray studies of *S.* Typhimurium regulons [100,111,114,141,142], as well as transcriptional profiling of *S.* Typhimurium 4/74 under infection-relevant *in vitro* growth conditions [32] and inside murine macrophages [41]. Detailed description of the pathway curation process and all relevant scripts can be found at https://github.com/apredeus/salmonella_pathways. Additionally, KEGG pathways for *S*. Typhimurium SL1344 were obtained using the KEGGREST R package v1.28.0, and gene ontology (GO) mapping of SL1344 genes was extracted from SL1344 protein annotation by InterProScan v5.41-78. See Dataset S2 for the entire dataset. The combined pathways were used to annotate gene sets up-regulated in the vacuole and cytosol using the clusterProfiler R package v3.16.1. The resulting overlaps were visualized using ggplot2 v3.3.2.

### Fluorescence microscopy

HeLa cells were seeded on acid-washed, 12 mm glass coverslips (#1.5 thickness, Fisher Scientific) in 24-well plates. Infected HeLa cells were fixed with 2.5% (w/v) paraformaldehyde in PBS for 10 min at 37°C. The immunostaining procedure has been described previously [22]. Rabbit anti-SiiE serum (kindly provided by Michael Hensel) was used at a dilution of 1:200. Digitonin permeabilization to deliver anti-*Salmonella* LPS antibodies (*Salmonella* O-Antiserum Group B Factors 1, 4, 5, 12; Difco; 1:300 dilution) directly to the cytosol was as described previously [14]. Cells were stained with Hoechst 33342 (1:10,000 in DDH_2_O, Life Technologies) to label DNA and coverslips were mounted onto glass slides using Mowiol. Samples were visualized on a Leica DM4000 upright fluorescence microscope for scoring the number of bacteria per cell, the proportion of cytosolic bacteria and quantification of transcriptional reporter activity. ImageJ software was used to quantify the activity of transcriptional reporters at the individual bacterium level. Images in the red (mCherry) and green (GFP) channels were acquired under 63x magnification on a Leica DM4000 upright fluorescent microscope using the same acquisition settings (exposure and gain) for each group of transcriptional reporters (determined by the reporter with the highest intensity in the green fluorescence channel). A minimum of 100 cytosolic and 100 vacuolar bacteria, from cells with ≥100 bacteria and 2-40 bacteria, respectively, acquired from ≥5 fields of view were quantified for each transcriptional reporter from each experiment. From grayscale images, well-defined, individual bacteria were randomly chosen on the mCherry channel (represents all bacteria) and manually outlined, converted to a binary image and the pixel intensity associated with each identified particle (i.e. one bacterium) quantified from the corresponding GFP channel image. GFP fluorescence/bacterium was subsequently plotted as the mean fluorescence intensity (MFI).

Image acquisition on a Leica SP8 Scanning Point confocal microscope was using the sequential acquisition mode through an optical section of 0.3 µm in the z-axis. Images are maximum intensity projections of z-stacks.

### Statistical analysis

All experiments were conducted on at least three separate occasions, unless otherwise indicated, and results are presented as mean ± SD. Except for the RNA-seq data (see separate paragraph on RNA-seq data analysis), statistical analyses were performed using one-way analysis of variance (ANOVA) with Dunnett’s post-hoc test or Student’s t-test (GraphPad Prism). A p-value of ≤0.05 was considered significant.

## Acknowledgements

We kindly thank Michael McClelland, Corrie Detweiler, Carsten Kroeger, Denise Monack, Martina Sassone-Corsi and Manuela Raffatellu for sharing bacterial mutants; Michael Hensel for providing the anti-SiiE antibody; Olivia Steele-Mortimer for providing the pCHAR-Duo plasmid; Zeus Saldana-Ahuactzi and Sadie Izaguirre for generating the P*zinT-gfpmut3* and P*mgtC-gfpmut3* constructs; Isabelle Hautefort for invaluable advice about bacterial RNA purification; Andrea Battistoni for helpful discussions about zinc transporters and Jean Celli and Johanna Elfenbein for critical reading of this manuscript.

## Supporting Information

**Dataset S1 (separate file).** RNA-seq data set. All genes/sRNAs with mapped reads were reported as TPM and a threshold TPM value of 10 was used as a cut-off to distinguish gene expression from background. Genes/sRNAs were deemed differentially expressed using fold-change criteria of ≥1.40 (cytosol up-regulated) and ≤0.71 (vacuole up-regulated) for WTM/EBSS treatment conditions (cells highlighted in yellow in Tab 1). Statistical significance was assessed using two independent platforms, Bayesian interference of transcripts from sequencing data (BitSeq) and rank products (RankProduct). For Bitseq analysis, we arbitrarily set a Probability of Positive Log Ratio (PPLR) of ≥0.90 for “up cytosol” genes/sRNAs and ≤0.1 for “up vacuole” genes/sRNAs for statistical significance (cells highlighted in green in Tab 1). For RankProduct analysis, the cutoff for differentially expressed genes was arbitrarily set to a percentage of false positive (pfp) value of ≤0.2 (cells highlighted in green in Tab 1) for statistical significance. Tab 2 is the short list of “up cytosol” genes/sRNAs that passed statistical significance. Tab 3 is the short-list of “up vacuole” genes/sRNAs that passed statistical significance. Expression profiles of these genes/sRNAs in 22 distinct infection-relevant *in vitro* growth conditions is shown in Tab 2 and Tab 3 (http://bioinf.gen.tcd.ie/cgi-bin/salcom.pl?db=salcom_mac_HL).

**Dataset 2 (separate file).** Custom pathway enrichment analysis of “up cytosol” and “up vacuole” genes/sRNAs. Tab 1 represents the curated list shown in Figure 2. Tab 2 shows the starting entire dataset.

---

**Figure S1.**
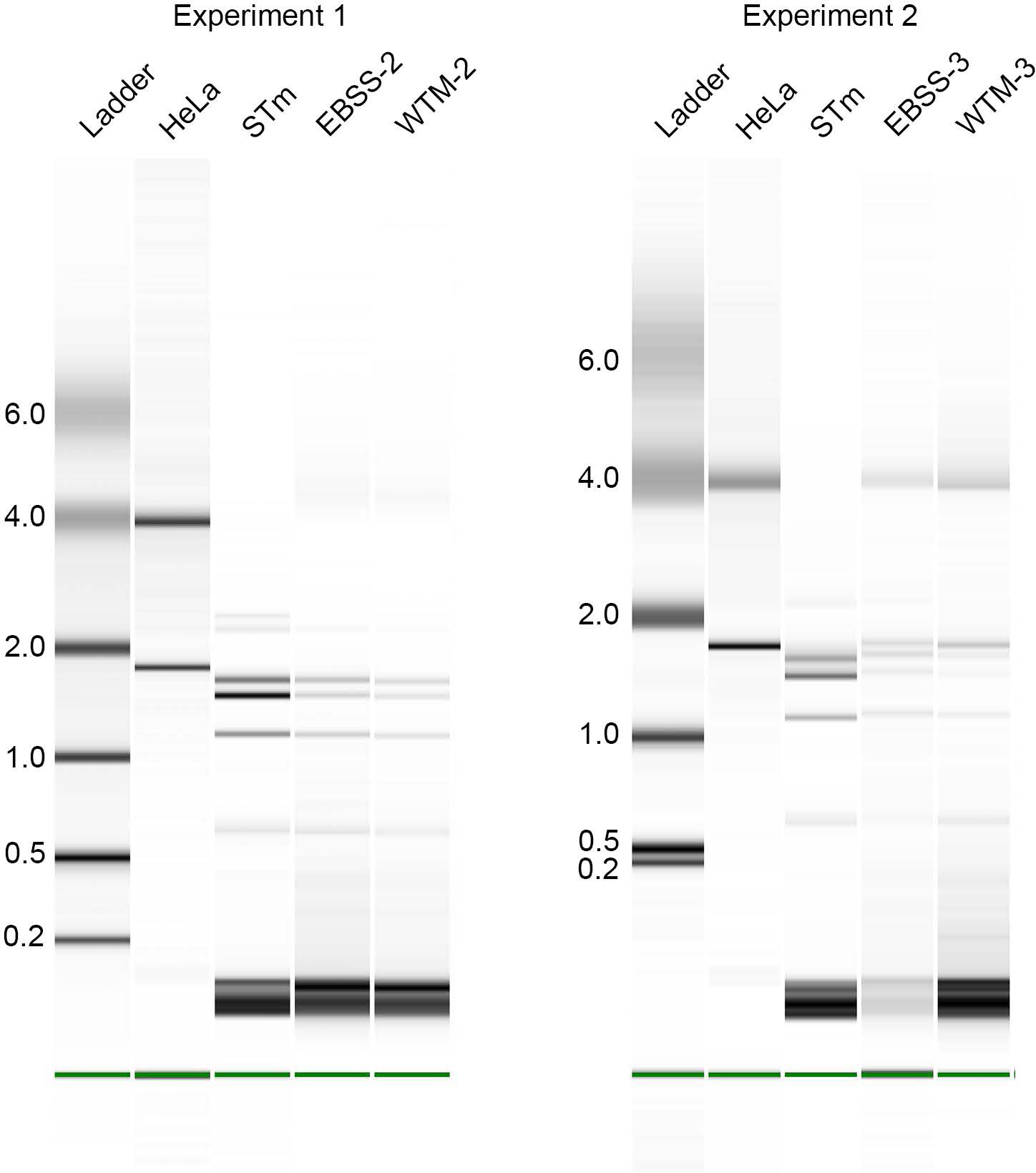
Enrichment of bacterial RNA from infected cells. Total RNA was extracted from HeLa epithelial cells (HeLa), wild-type S. Typhimurium SL1344 grown to late log-phase (STm), S. Typhimurium isolated from Earle’s balance salt solution (EBSS)-treated cells at 8 h p.i. or *S.* Typhimurium isolated from wortmannin (WTM)-treated cells at 8 h p.i. RNA quality was analyzed by electrophoretic separation using an Agilent Bioanalyzer 2100. Ladder sizes shown in kb.

**Figure S2.**
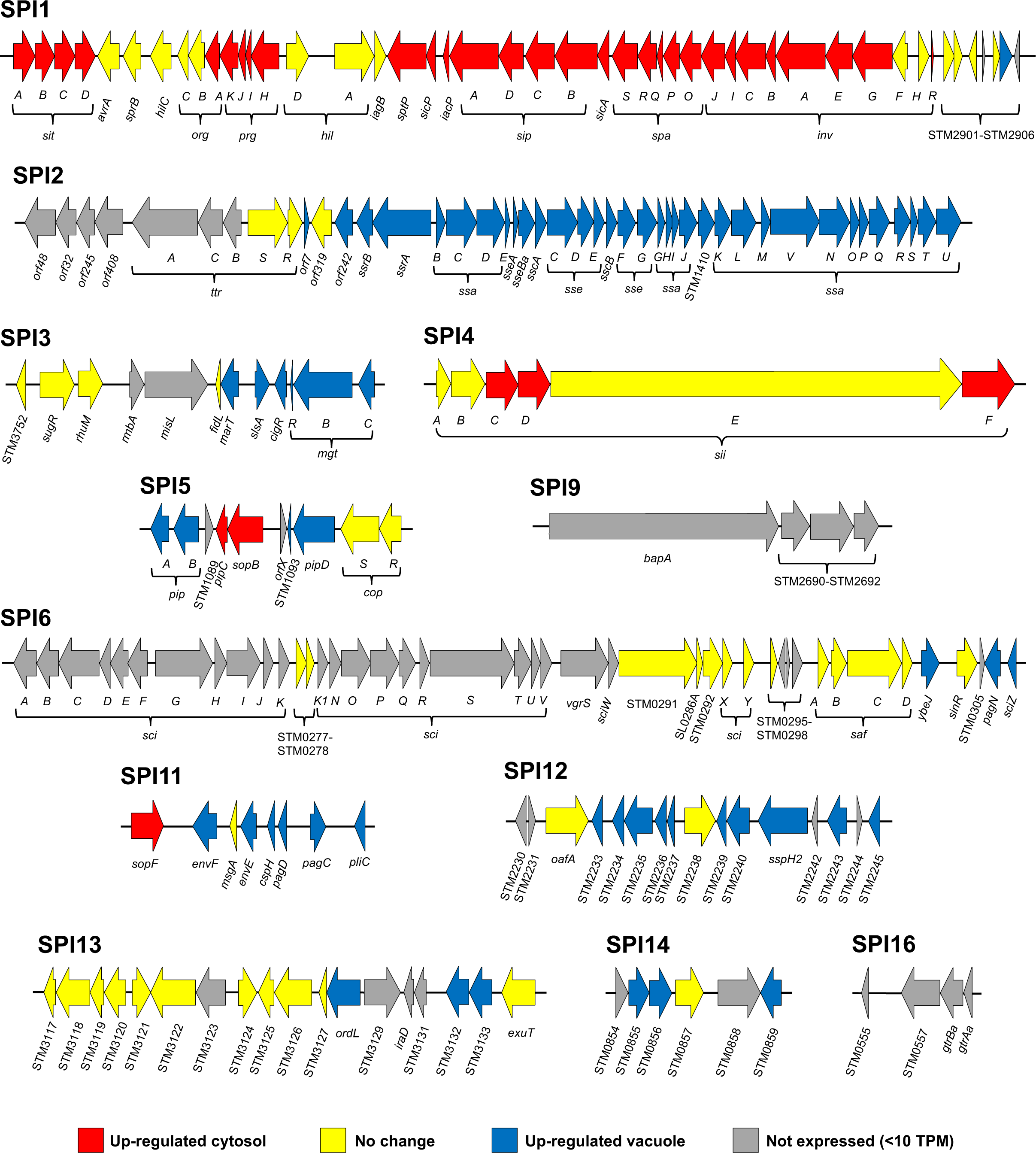
Relative expression of the different PAIs of *S*. Typhimurium. Each arrow represents an individual gene to scale within each PAI. The different islands are also scaled against each other. The color of each arrow represents relative gene expression – red arrows depict genes up-regulated in the cytosol (≥1.40-fold change WTM/EBSS), blue are genes up-regulated in the vacuole (≤0.71-fold change WTM/EBSS), yellow are genes with unchanged expression (0.72-1.39-fold change) and grey arrows are genes with a TPM value <10 and considered not expressed. See Dataset S1 for the entire data set. Adapted from Srikumar et al. (41)

**Figure S3.**
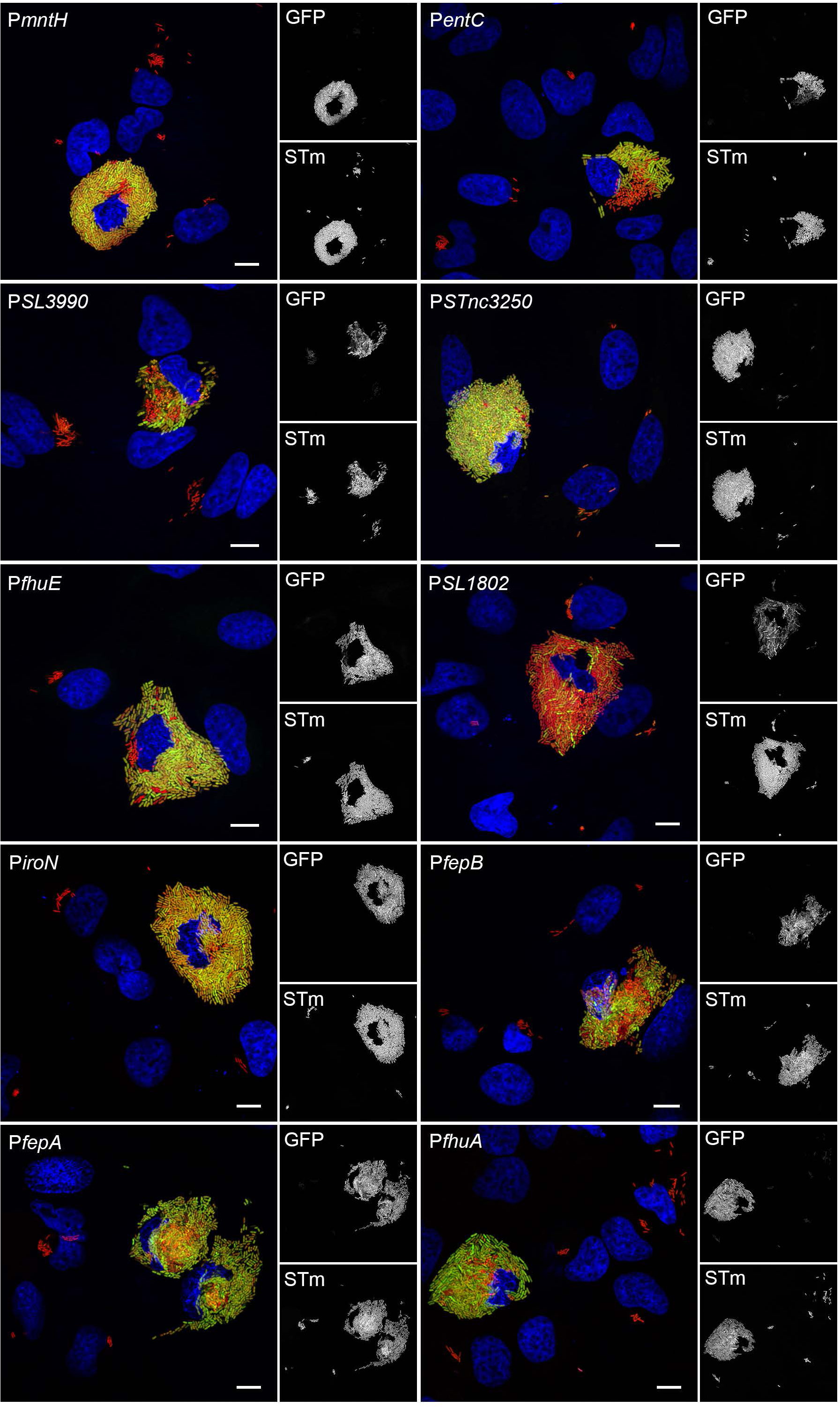
Additional iron-associated genes/sRNAs that are cytosol-specific. Epithelial cells were infected with mCherry-S. Typhimurium harboring *gfpmut3* transcriptional reporters. At 8 h p.i., cells were fixed and stained with Hoechst 33342 to detect DNA Representative confocal microscopy images show induction of *mntH, entC,* SL3990, STnc3250, *fhuE,* SL1802, *iroN, fepB, fepA* and *fhuA* promoters in cytosolic bacteria. Green= transcriptional reporter, red= S. Typhimurium, blue= DNA. Scale bars are 10 µm.

**Figure S4.**
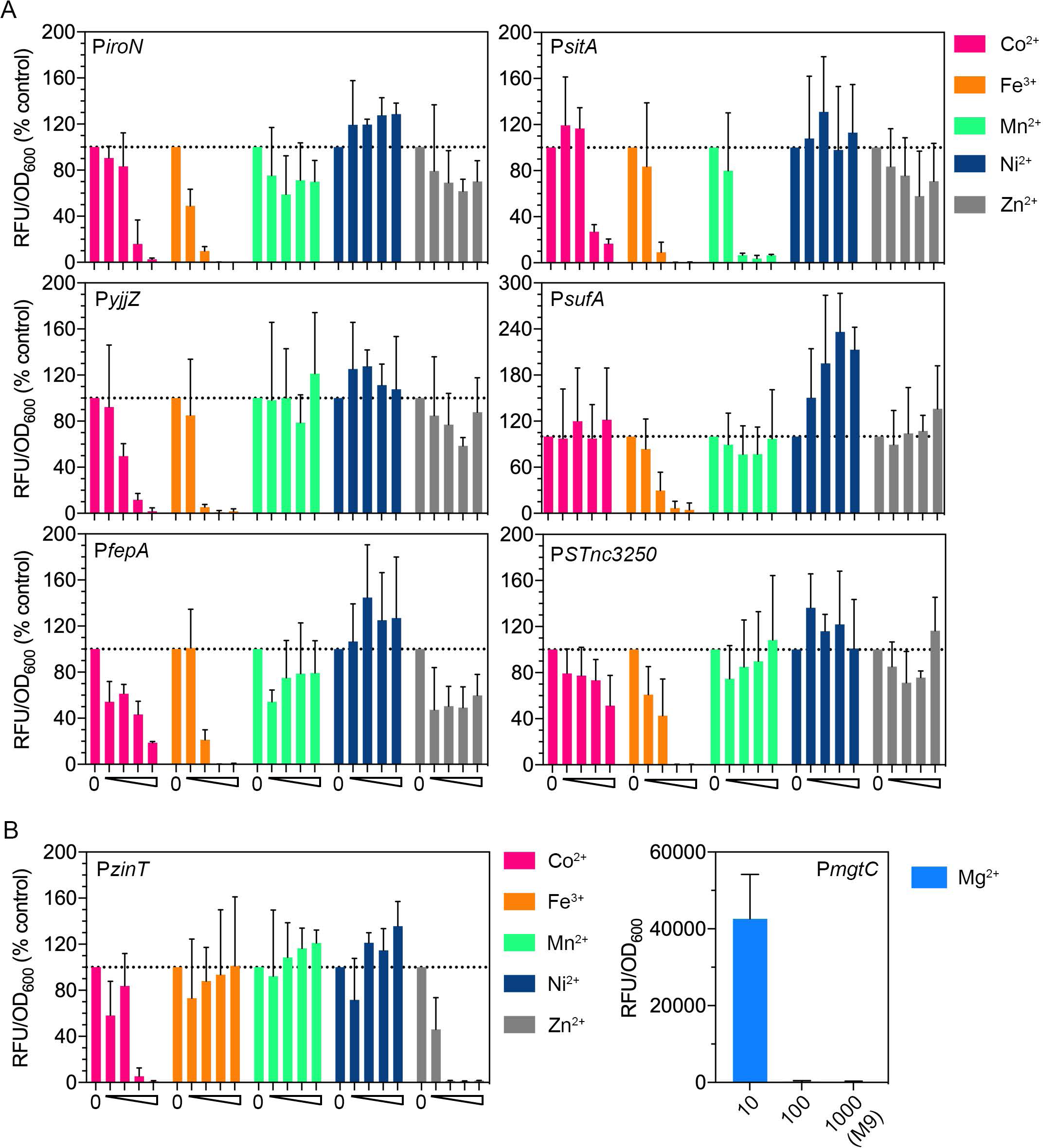
The common metal regulator of cytosol-specific gene/sRNA transcription is iron. **(A).** GFP fluorescence in mCherry-S. Typhimurium harboring *PiroN-gfpmut3, PsitA-gfpmut3, PyjjZ-gfpmut3, PsufA-gfpmut3, PfepA-gfpmut3* or *PSTnc3250-gfpmut3* reporter plasmids. Bacteria were grown shaking overnight at 37^°^C for 16 h in M9-supplemented media containing increasing concentrations of CoCl_2_, FeCl_3_, MnCl_2_, NiCl_2_ or ZnCl_2_ (0.1 µM, 1 µM, 10 µM or 100 µM). No added cation (0) served as the control. The relative fluorescence units (RFU) were normalized to 00_600_ and expressed as a percentage of control. Background GFP fluorescence of mCherry-S. Typhimurium (no reporter) was subtracted from all values. n≥3 independent experiments. (B) GFP fluorescence in mCherry-S. Typhimurium harboring *PzinT-gfpmut3* or *PmgtC-gfpmut3* plasmids. Growth and analysis of zinTexpression was as described in (A).Standard M9 minimal media contains 1 mM MgS0_4_ which completely represses *mgtC* expression. The *effect* of Mg^2+^ concentration on *mgtC* expression was therefore assessed in M9-supplemented media containing decreasing amounts of MgS0_4_ i.e. 1000 µM (standard), 100 µM and 10 µM. Mg^2+^ concentrations lower than 10 µM considerably impacted bacterial growth. n=3 independent experiments.

**Figure S5.**
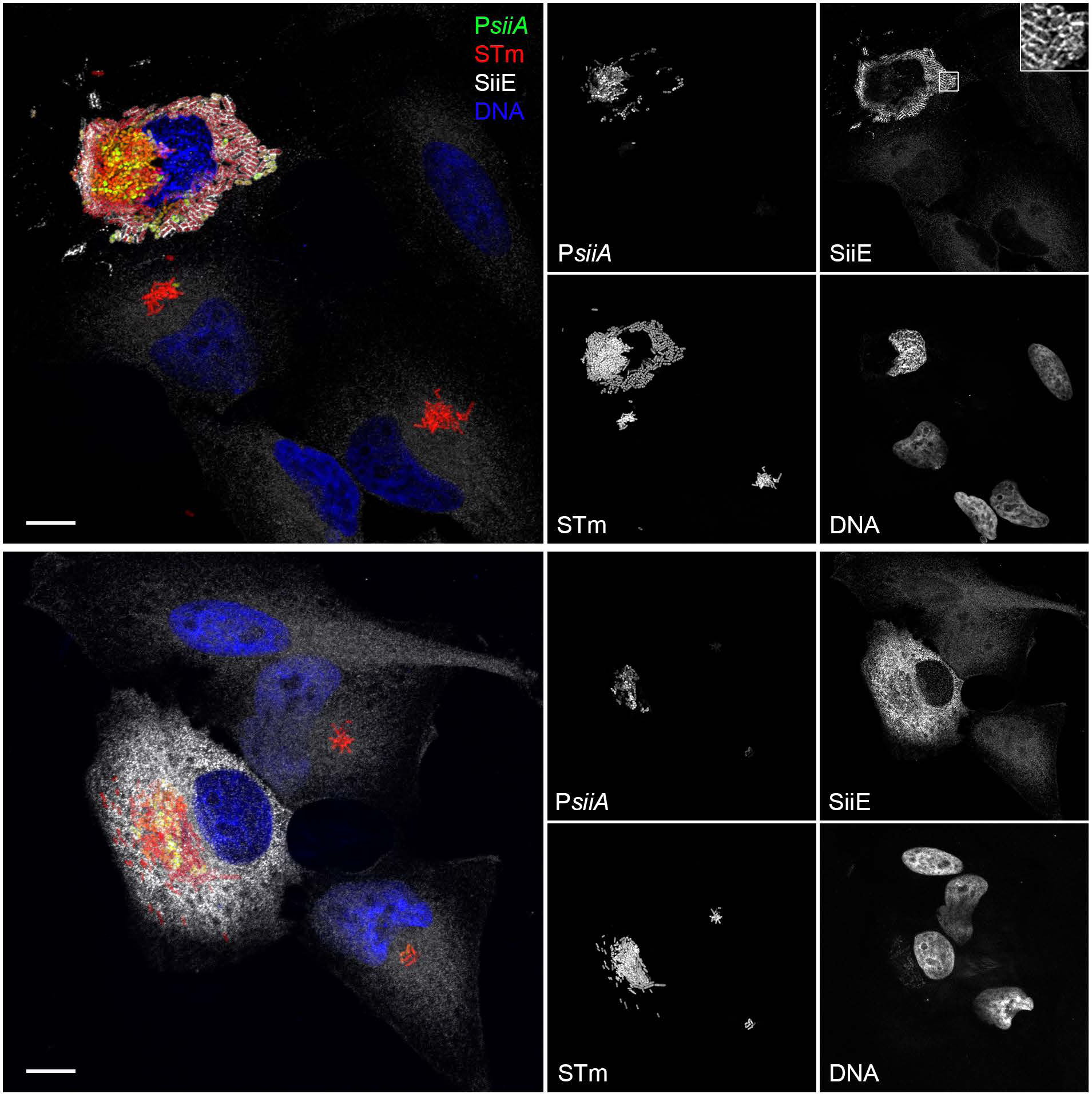
Cytosolic bacteria produce SiiE. Epithelial cells seeded on coverslips were infected with mCherry-S. Typhimurium harboring a *PsiiA-gfpmut3* transcriptional reporter. At 8 h p.i., cells were fixed and immunostained with polyclonal antibodies directed against SiiE. DNA was stained with Hoechst 33342. Representative confocal microscopy images show SiiE attached to (upper panel) or secreted by (lower panel) cytosolic bacteria. Inset shows enlargement of boxed area. Green= *PsiiA-gfpmut3* reporter, red= S. Typhimurium, white= SiiE, blue= DNA. Scale bars are 10 µm.

**Figure S6.**
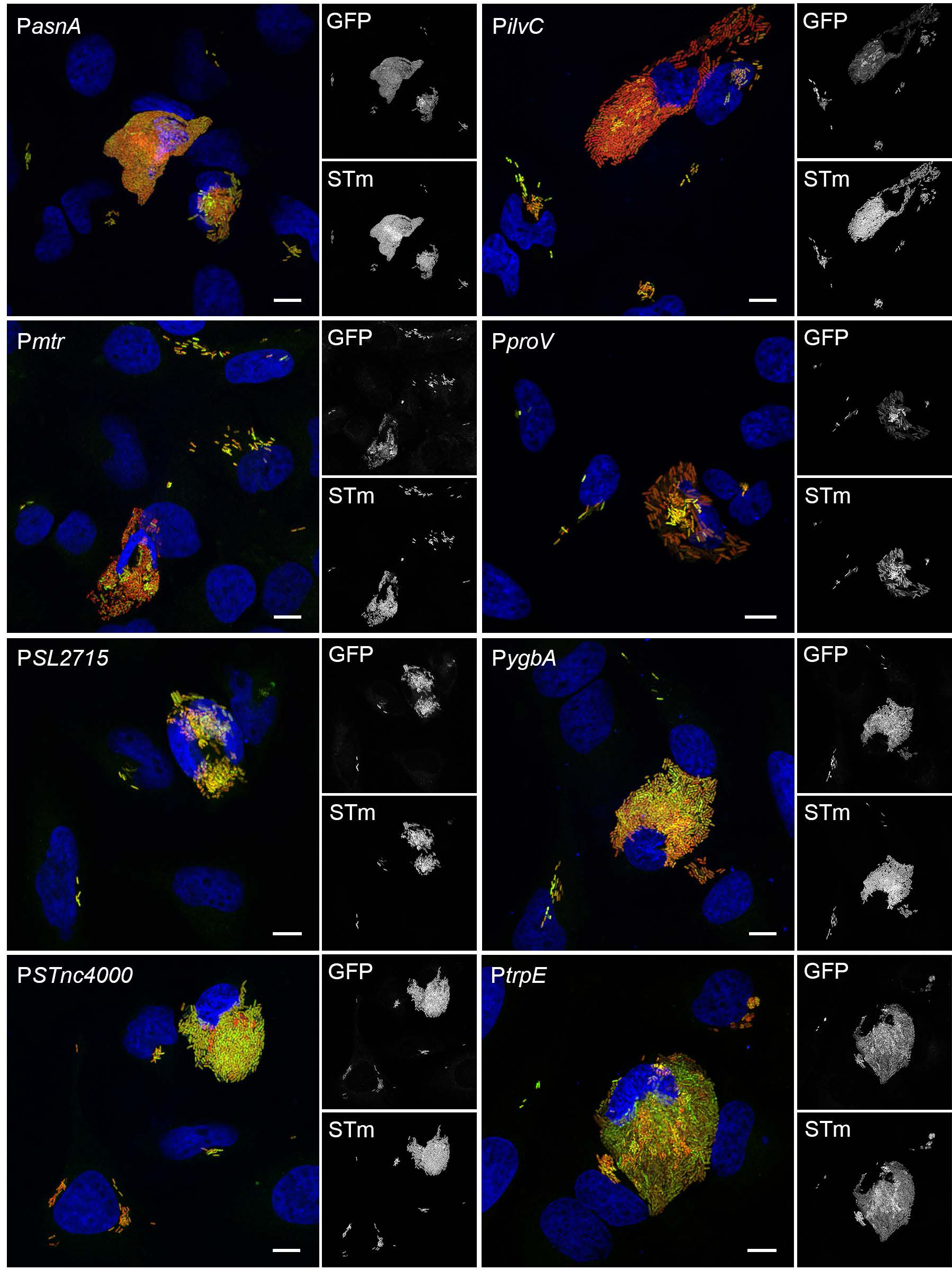
RNA-seq-predicted genes/sRNAs that were not confirmed by transcriptional reporters. Epithelial cells seeded on coverslips were infected with mCherry-S. Typhimurium harboring *gfpmut3* transcriptional reporters. At 8 h p.i., cells were fixed and stained with Hoechst 33342 to detect DNA. Representative confocal microscopy images show equivalent expression of *asnA, ilvC, mfr, proV,* SL1344_2715, *ygbA,* STnc4000 and *trpE* promoters in vacuolar and cytosolic bacteria. Green = transcriptional reporter, red = S. Typhimurium, blue = DNA. Scale bars are 10 µm.

**Figure S7.**
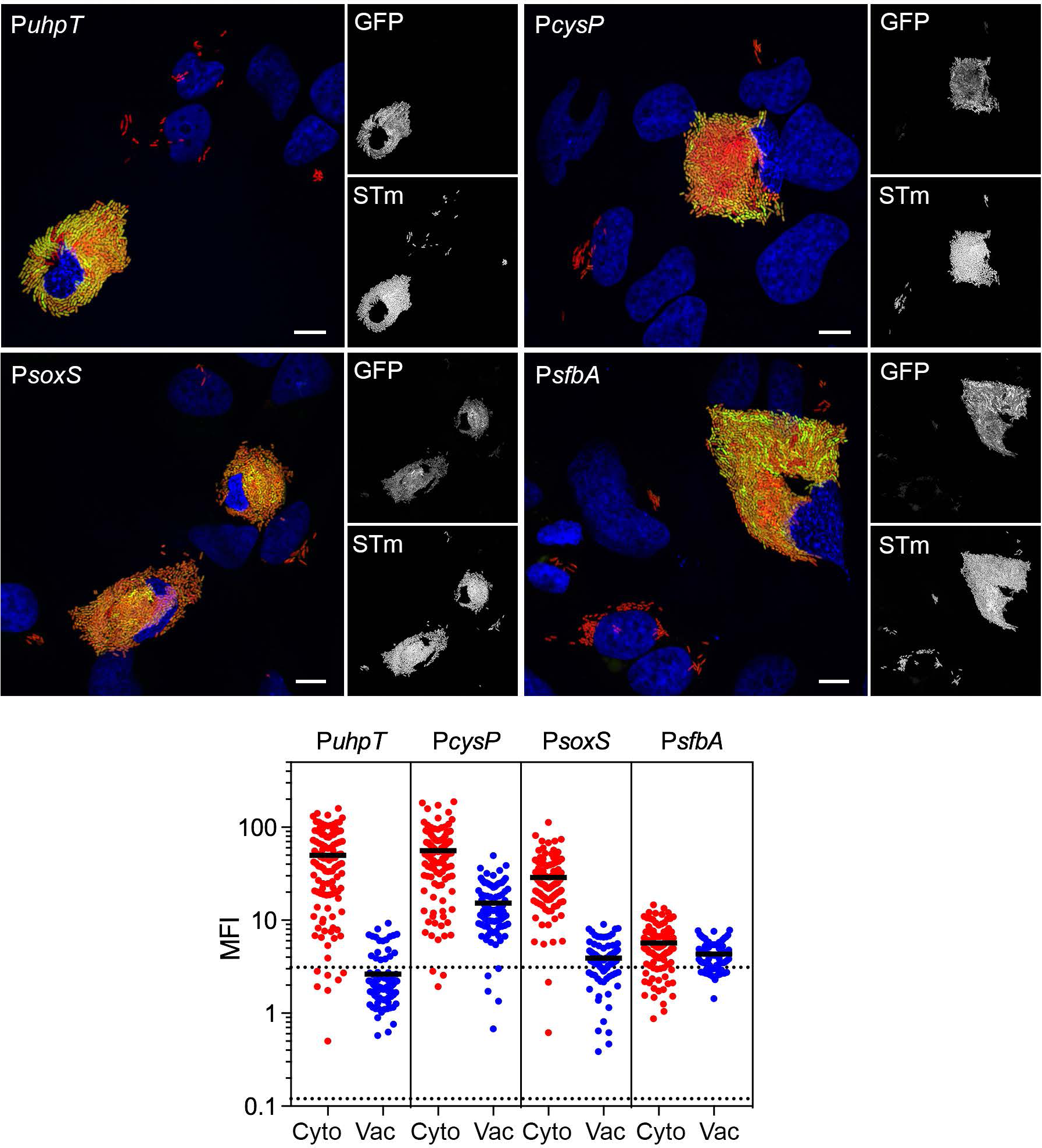
Identification of additional *S.* Typhimurium genes that are up-regulated in the epithelial cytosol. Upper panels: Epithelial cells seeded on coverslips were infected with mCherry-S. Typhimurium harboring *gfpmut3* transcriptional reporters. At 8 h p.i., cells were fixed and stained with Hoechst 33342 to detect DNA. Representative confocal microscopy images show induction of *uhpT, cysP, soxS* and *sfbA* promoters in cytosolic bacteria. Green = transcriptional reporter, red = S. Typhimurium, blue = DNA. Scale bars are 10 µm. Lower panel: Quantification of the MFI of GFP signal in individual bacteria. Bacteria were designated as being cytosolic (Cyto) or vacuolar (Vac) if residing within cells with ≥100 bacteria or 2-40 bacteria, respectively. Each dot represents one bacterium; the solid black line indicates the mean. Acquisition parameters (acquisition time and gain) were set-up using *PcysP-gfpmut3* (the highest GFP signal intensity) and these same parameters were applied throughout. The MFI for each bacterium was determine using lmageJ. Dashed lines indicate the range of background fluorescence in the GFP channel measured for mCherry-S. Typhimurium (no reporter). n=2 independent experiments.

